# Region-specific KCC2 rescue by rhIGF-1 and oxytocin in a mouse model of Rett syndrome

**DOI:** 10.1101/2021.09.25.460342

**Authors:** Valentina Gigliucci, Jasper Teutsch, Marc Woodbury-Smith, Mirko Luoni, Marta Busnelli, Bice Chini, Abhishek Banerjee

## Abstract

Rett syndrome (RTT) is characterized by dysfunction in neuronal excitation/inhibition (E/I) balance, potentially impacting seizure susceptibility via deficits in K^+^/Cl^-^ cotransporter 2 (KCC2) function. Mice lacking the Methyl-CpG binding protein 2 (MeCP2) recapitulate many symptoms of RTT, and recombinant human insulin-like growth factor-1 (rhIGF-1) restores KCC2 expression and E/I balance in MeCP2 KO mice. However, clinical trial outcomes of rhIGF-1 in RTT have been variable, and increasing its therapeutic efficacy is highly desirable. To this end, the neuropeptide oxytocin (OXT) is promising, as it also critically modulates KCC2 function during early postnatal development. We measured basal KCC2 expression levels in MeCP2 KO mice and identified three key frontal brain regions showing KCC2 alterations in young adult mice but not in postnatal P10 animals. We thus hypothesized that deficits in an IGF-1/OXT signaling crosstalk modulating KCC2 may occur in RTT during postnatal development. Consistently, we detected alterations of IGF-1 receptor (IGF-1R) and OXT receptor (OXTR) levels in those brain areas. rhIGF-1 and OXT treatments in KO mice rescued KCC2 expression in a region-specific and complementary manner. These results suggest that region-selective combinatorial pharmacotherapeutic strategies could be the most effective at normalizing E/I balance in key brain regions subtending the RTT pathophysiology.

## Introduction

Rett syndrome (RTT) is a pervasive neurodevelopmental disorder arising from loss-of-function mutations in the methyl-CpG binding protein 2 (*Mecp2*) gene (Banerjee et al. 2019). In the brain, the lack of MeCP2 protein results in an imbalance in neuronal and synaptic excitation and inhibition (E/I), potentially leading to susceptibility to seizure frequently seen in RTT patients (Glaze et al. 2010). K^+^/Cl^-^ co-transporter 2 (KCC2) plays an important role in the establishment and maintenance of appropriate GABAergic tone and E/I balance during early development. KCC2 levels are very low prenatally, but increase around birth resulting in changes in the excitatory-to-inhibitory polarity in GABAergic neurotransmission (Ben-Ari et al. 2012). Pathogenic variants of the human KCC2 gene, *SLC12A5*, have been implicated in both epilepsy and autism spectrum disorder (ASD) (Stödberg et al. 2015; Krupp et al. 2017), and recent reports in animal models highlight the importance of KCC2 in early development and the long-term effects of its dysregulation. Resetting a correct E/I balance during development may offer a new therapeutic strategy in RTT and related neurodevelopmental disorders, and KCC2 represents a promising therapeutic target (Banerjee et al. 2016; Tang et al. 2019). KCC2 levels are altered in the brain of RTT patients (Duarte et al. 2013; Hinz et al. 2019) and also in MeCP2 KO mice (Banerjee et al. 2016), a valuable model to study the pathophysiology of RTT, recapitulating many of its symptoms (Banerjee et al. 2012). Hence, endogenous and exogenous factors capable of impacting KCC2 expression and function in RTT are worthy of careful investigation.

Two critical candidate regulators of KCC2 function are insulin-like growth factor (IGF-1) and the neuropeptide oxytocin (OXT). IGF-1 is a growth factor crucial for neuronal survival and maturation and it has been known to modulate KCC2 activity and accelerate neuronal maturation by promoting the GABA shift. Administration of recombinant full-length human IGF-1 (rhIGF-1) to MeCP2 KO mice resulted in the rescue of both KCC2 expression and synaptic transmission deficits (Banerjee et al., 2016). Clinical trials in RTT girls showed that rhIGF-1 treatment ameliorated some symptoms, such as anxiety and respiratory abnormalities; however, the clinical outcomes have been variable (Khwaja et al. 2014; Pini et al. 2016; O’Leary et al. 2018). OXT also has a crucial role in neurodevelopment by impacting the correct timing of the GABA shift (Tyzio et al. 2014), and it directly modulates KCC2 expression and function (Leonzino et al. 2016). To our knowledge, OXT has never been experimented on RTT patients, however it is currently under clinical investigation as a potential treatment for ASD (DeMayo et al. 2017; Baker and Stavropoulos 2020), making it a relevant therapeutic candidate for RTT.

As the importance of KCC2 in neurodevelopment, intellectual disability (ID) and developmental disorders, including RTT, is poorly characterized in humans, we first analyzed the co-expression network for the gene coding for KCC2, *SLC12A5*, in developing human brains, hypothesizing that its interacting partners would be key genes implicated in ASD, epilepsy, and ID. Having confirmed the potential relevance of KCC2 in neurodevelopmental disorders, we next moved to analyzing MeCP2 KO RTT mice. We measured regional KCC2 expression and found that KCC2 levels were selectively and specifically reduced in frontal brain areas involved in complex cognition, decision making, and social behaviour. Furthermore, we found that in regions with KCC2 deficiencies, IGF-1 receptor (IGF-1R) and OXT receptor (OXTR) levels were also altered. Since both IGF-1 and OXT signalling pathways converge on modulating KCC2, we hypothesized that these two factors cooperate to restore KCC2 levels in MeCP2 KO mice. Finally, treatment with rhIGF-1 or OXT showed region-specific non-overlapping rescue of KCC2 expression. These results suggest that OXT together with rhIGF-1 hold a therapeutic potential in RTT.

## Materials and Methods

### Animals

Experimental procedures were carried out in accordance with the guidelines of the Federal Veterinary Office of Switzerland and the Italian Ministry of Health according to the European Communities Council Directive 2010/63/EU. Mecp2^-/y^ hemizygous KO (MeCP2 KO) mice and WT littermates were obtained by breeding Mecp2^+/-^ heterozygous females with WT male mice. All mice belonged to the C57BL6/J strain. Mice were grouped with their WT siblings and housed at 24°C and variable humidity in a 12-h dark/light cycle (7:00 a.m. to 7:00 p.m.). For P10 animals, all siblings were kept with their mothers since birth and removed from the cages just before sacrifice. In this study, we used Mecp2 hemizygous male mice because Mecp2 heterozygous females display a delayed phenotype than males; indeed, hemizygous MeCP2 KO males display profound severity in various autonomic, sensory/motor, and cognitive phenotypes starting early in postnatal life and they are considered to effectively model the human disorder (Guy et al. 2001; Samaco et al. 2013; Castro et al. 2014; Banerjee et al. 2012).

### Drug administration

#### A) Systemic administration of rhIGF-1

Animals were weighed and injected i.p. once daily for 14 days starting at P30 with either vehicle (saline) or rhIGF-1 (2.5 mg/kg; Peprotech, USA) dissolved in saline with 0.01% Bovine Serum Albumin (weight/volume).

#### B) Intranasal OXT administration

OXT (0.3 IU/5 μL; Bachem, Switzerland) was administered intranasally, as previously described (Huang et al. 2014). For OXT, intranasal route was chosen to overcome the short half-life of OXT in blood circulation, and because intranasal administration is the preferential route in clinical trials in autistic subjects. For each treatment, mice received 0.3 IU in a volume of 5 μL. An amount of 1 IU of the solution contained 1.667 mg of synthetic OXT. Thus, 0.3 IU corresponded to 0.5001 mg of OXT for each administration of 5 μL. Mice were administered with OXT twice a day, once in the morning and once in the afternoon, for 10 days. Previous studies (Huang et al. 2014) reported that 7 consecutive days of OXT treatment is sufficient to determine an effect on social behavior and on OXTR levels.

All animals were euthanized 2 hours after the last treatment (P40 for OXT-treated mice, P44 for rhIGF-1-treated mice), brains were quickly removed from the skull and snap frozen at −25°C in isopentane (2-Methylbutane, Merck Life Science, Italy). Once frozen, coronal brain sections of 14 μm thickness were sliced with a cryostat, thaw-mounted on microscope slides pre-coated with chrome-alum-gelatin and kept at −80°C until further use.

### KCC2 and IGF-1R infrared immunofluorescence

On the day of immunostaining experiment, sections were defrosted, fixed with 0.2% paraformaldehyde in 0.1 M phosphate-buffered saline (pH 7.4), blocked with 5% Bovine Serum Albumin for 2 hours at room temperature, then incubated overnight at 4°C with anti-KCC2 (1:200; Merck Life Sciences, Italy, #07-432) or anti-IGF-1R β (1:100; Cell Signaling Technology, Italy, #9750). For nonspecific binding, no primary antibody was added to the incubation solution. Sections were then incubated with goat-anti-rabbit-IR800 secondary antibody (1:2000; Li-Cor Biosciences GmbH, Germany, #926-32211) and scanned using an Odyssey CLx scanner (Li-Cor Biosciences GmbH, Germany). Scanning settings were - laser: 0.5 for KCC2 staining and 2 for IGF-1R staining; channel: 800; quality: highest; resolution: 21 μm. Scans were exported and analyzed with the ImageStudio Lite v.5.2 software (Li-Cor Biosciences GmbH, Germany). Regions of interest (ROIs) were hand-drawn according to the reference brain atlas (Franklin and Paxinos 2007), non-specific binding values were subtracted, and the net fluorescence intensity value was divided by the ROI’s size to obtain final signal/pixel data. Labelling with IR Dyes implies that the signal measured should be truly representative of the protein abundance. The fluorescent tag is directly conjugated to the secondary antibody, and it has previously been demonstrated that this produces a truly linear readout when applied to tissue sections and tissue sample homogenates by Western blotting (Eaton et al. 2016). For each brain region, 2-3 coronal planes were analyzed, depending on the rostro-caudal extension, (MOB +5.00/+3.92 mm from Bregma, AON +3.20/+2.68 mm from Bregma, AONm +2.34/+2.22 mm from Bregma, aPIR +2.46/+1.98 mm from Bregma, PFC +3.20/+2.58 mm from Bregma, BLA −0.94/-1.34 mm from Bregma, Hipp CA2/3 −1.46/-2.18 mm from Bregma, according to Franklin and Paxinos 2007), 2-3 sections per coronal plane. Left and right hemispheres were initially analyzed separately and checked for any significant interhemispheric differences. Given that no statistical significance was found (**Supplementary Tables 1** and **2**), all data were pooled together for the group analysis.

### OXTR autoradiography

On the day of autoradiography experiment, sections were defrosted, fixed with 0.2% paraformaldehyde in 0.1 M phosphate-buffered saline (pH 7.4), rinsed with 0.1% Bovine Serum Albumin in 50 mM Tris–HCl buffer (pH 7.4), and incubated 2 hours at room temperature with a 0.02 nM solution of the radio-iodinated OXTR antagonist ornithine vasotocin analogue ([^125^I]-OVTA, specific activity 2200 Ci/mmol; Perkin Elmer, MA, USA in 50 mM Tris–HCl, 0.025% bacitracin, 5 mM MgCl_2_, 0.1% Bovine Serum Albumin). Sections immediately adjacent to the ones used for [^125^I]-OVTA binding were used to determine non-specific binding by addition of 2 μM OXT to the incubation solution. Excessive ligand was washed out, and the slides were quickly dried and exposed to Biomax MR Films (Carestream, USA, #891-2560) for 72 hours. The final autoradiograms were digitalized by grayscale high-resolution scanning (600×600 dpi) and analysis of the optical binding density of the brain ROIs was carried out using the ImageJ 1.47v software (NIH, USA). ROIs were identified by comparison with a reference mouse brain atlas (Franklin and Paxinos 2007) and manually delineated with the ROI manager tool of the software. Specific densitometric grey intensity was calculated by subtraction of the grey level of the respective section treated for non-specific binding. For each brain region, 2-3 coronal planes were analyzed, depending on the rostro-caudal extension, (MOB +5.00/+3.92 mm from Bregma, AON +3.20/+2.68 mm from Bregma, AONm +2.34/+2.22 mm from Bregma, aPIR +2.46/+1.98 mm from Bregma, PFC +3.20/+2.58 mm from Bregma, BLA −0.94/-1.34 mm from Bregma, Hipp CA2/3−1.46/-2.18 mm from Bregma, according to Franklin and Paxinos 2007), 2-3 sections per coronal plane. Left and right hemispheres were initially analyzed separately and checked for any significant interhemispheric differences. Given that no statistical significance was found (**Supplementary Tables 1** and **2**), all data were pooled together for the group analysis.

### Statistical analysis

Data was analyzed with the GraphPad Prism software (v. 6.1, GraphPad Software, Inc., USA). For the direct comparisons of two groups, unpaired, double-tailed t-test was used. The Sidak-Bonferroni correction for multiple t-testing was applied (alpha = 5.0%). When SDs of the compared groups were significantly different, the Welch’s correction was applied. When data distribution was not normal, a non-parametric Mann Whitney U-test was used. Pharmacological experiments were analyzed by one-way ANOVA followed by Holm-Sidak’s multiple comparisons *post-hoc* test. Results were deemed significant when p < 0.05.

## Results

### Lack of MeCP2 leads to region-specific alterations in KCC2 expression

MeCP2 expression increases during development and is believed to parallel synaptogenesis (Feldman et al. 2016). During development, KCC2 is transcriptionally modulated by MeCP2 (Tang et al. 2016), suggesting its involvement in RTT. Our analysis of the co-expression modules of the KCC2 coding gene (*SLC12A5*) corroborated our hypothesis. Indeed, *SLC12A5* is part of a synaptic gene network associated with ASD, ID, and epilepsy, the three neurological conditions most strongly associated with RTT (**Supplementary Data** and **Supplementary Fig. 1**). The demonstration that in the KCC2 (*SLC12A5*) gene coexpression network there is an over-representation for ASD, ID, and epilepsy candidate genes strengthens the hypothesis that KCC2 and its interacting partners are important across these disorders, and therefore represent potential druggable targets (Pozzi and Chini, 2021).

We further hypothesized that in the male MeCP2 KO mice, when most of the synaptogenesis events have already taken place, KCC2 levels should be altered. To test this, we used P40-44 MeCP2 KO mice and determined total KCC2 levels by infrared immunofluorescence in distributed brain regions related to RTT, with particular focus on areas involved in processing social stimuli and social behavior (**Fig. 1A** and **Supplementary Fig. 2**). We selected several areas of the olfactory system (the main olfactory bulb - MOB, anterior olfactory nucleus - AON, medial part of the anterior olfactory nucleus – AONm, and the anterior part of the piriform cortex – aPIR), as olfactory function and circuitry maturation is altered by the lack of MeCP2 (Degano et al. 2014; Martínez-Rodríguez et al. 2019). The olfactory system is fundamental for social behavior in rodents, and all the aforementioned regions express high levels of OXTR (Gigliucci et al. 2014; Huang et al. 2014). Next, we focused on the basolateral amygdala (BLA), one of the projection targets of the aPIR, which is involved in social learning and a potential key area for the treatment of motor and social symptoms in RTT (Hsu et al. 2020). We also looked at the prefrontal cortex (PFC), as PFC’s E/I balance is known to be altered in MeCP2 KO mice, and selective intervention in this area can rescue the cognitive impairments seen in these mice (Sceniak et al. 2016; Howell et al. 2017; Banerjee et al. 2020). Finally, hippocampal deficits have been repeatedly reported in RTT mice models (extensively reviewed by Li et al. 2019), therefore we extended our analysis to the hippocampus. We focused particularly on the hippocampal CA2/3 subfield (Hipp CA2/3), as OXTR labelling is concentrated in this subregion (Gigliucci et al. 2014). We detected and quantified KCC2 in all of the aforementioned regions.

**Fig.1.**
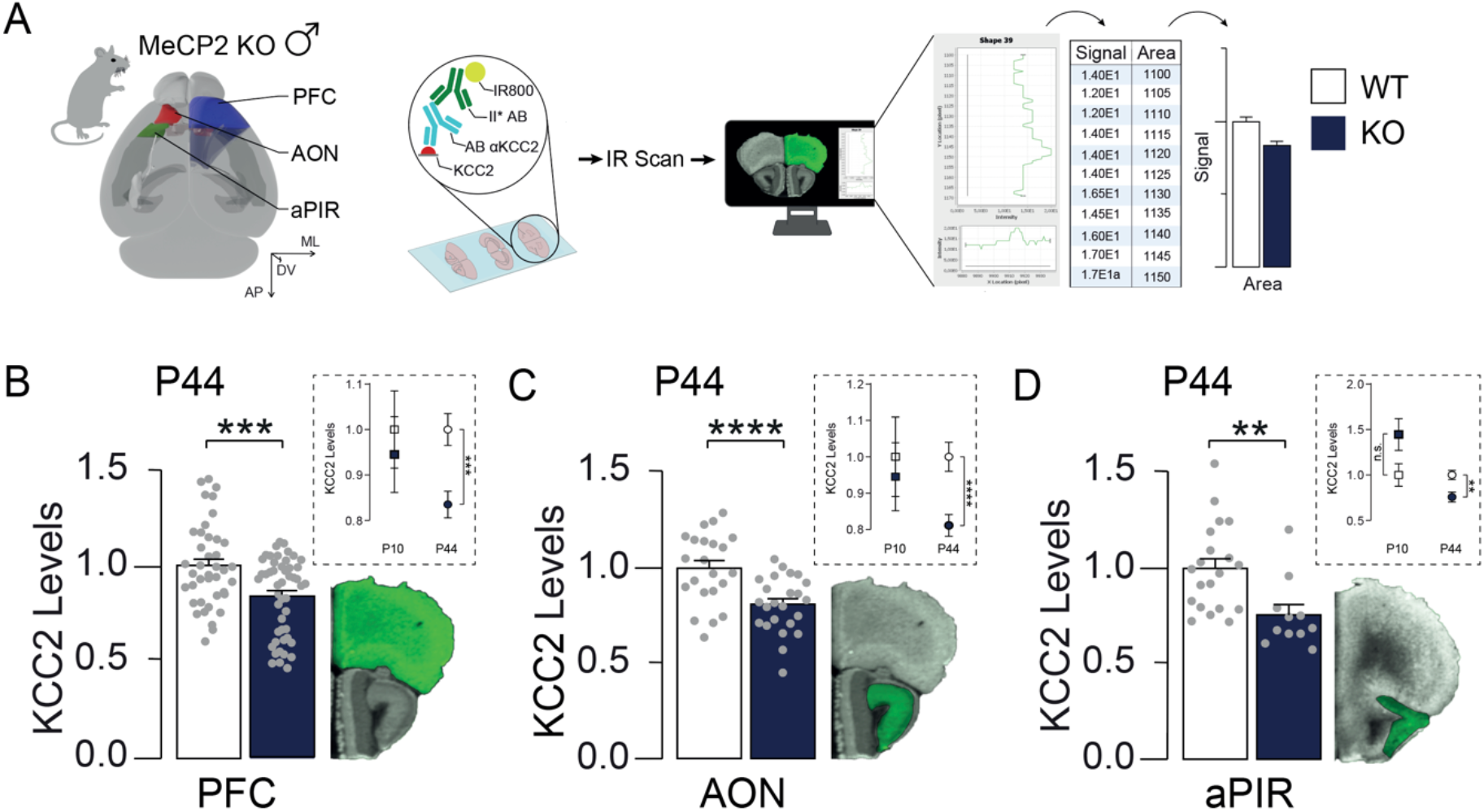
MeCP2 KO mice display region-specific alterations in total KCC2 levels. Lack of MeCP2 results in variable reductions of KCC2 levels across the brain regions analyzed. (**A**) Reconstruction of a mouse brain displaying the location of the areas analyzed for KCC2 expression, with a schematic representation of the infrared immunofluorescence procedure and analysis that was carried out; (**B-C-D**) Bar graphs of KCC2 levels (plotted as fold change over WT-Veh) showing that KCC2 level was significantly reduced selectively in the (C) Prefrontal Cortex (PFC), (D) Anterior Olfactory Nucleus (AON) and (E) Piriform Cortex, anterior part (aPIR) in P44 MeCP2 KO mice in comparison to WT-Veh controls. Such reductions were not present in these areas at an earlier developmental time point (P10) (*inset*). On the side, examples of infrared KCC2 labelled coronal sections. Areas in green represent the regions of interest quantified in the analysis. PFC = blue, AON = red, aPIR = green. Data expressed as mean ± SEM, single dots represent single observations from 3-4 animals per group, 2 or 3 coronal planes per brain region, 2-3 sections per coronal plane, left and right hemisphere data pooled together. **p < 0.01, ***p < 0.001, ****p < 0.0001 vs WT-Veh.

Our data demonstrates that, in MeCP2 KO mice, alterations in KCC2 expression are regionspecific. In particular, the PFC (t_(87)_ = 3.643; p < 0.001), AON (t_(44)_ = 3.836; p<0.001) and aPIR (t_(29)_ = 3.041; p < 0.01) showed significantly reduced KCC2 expression levels in MeCP2 KO mice when compared to WT littermates (**Fig. 1B-C-D;** all corrected for multiple comparisons). In contrast, the other regions analyzed (MOB, AONm, BLA and Hipp CA2/3) did not show alterations in KCC2 expression in MeCP2 KO mice in comparison with WT littermates (**Supplementary Fig. 2B-C-D-E**). These alterations of KCC2 levels in the PFC, AON, and the aPIR were not detected at an early stage of post-natal development (P10; **Fig. 1 C-D-E** *inset*). This indicates that the effects of the lack of MeCP2 arise later during postnatal development (after P10), and do not impact on adult total KCC2 levels uniformly across brain regions, highlighting some key areas that could be targeted further in our experiments.

### KCC2 alterations are accompanied with changes in OXTR and IGF-1R expression

We next assessed whether the regional reductions in KCC2 levels in MeCP2 KO mice may be linked to variations in the expression levels of IGF-1R and OXTR, the two receptors involved in IGF-1 and OXT signalling and KCC2 regulation. To investigate this, IGF-1R levels were quantified using infrared immunofluorescence and OXTR levels were quantified using radioligand autoradiography in MeCP2 KO mice and compared with WT littermates.

After correction for multiple comparisons, we found that in MeCP2 KO mice, IGF-1R and OXTR levels were altered in the three key brain areas in which KCC2 levels were reduced - the PFC, AON and the aPIR. In particular, IGF-1R levels were increased in the PFC (Welch-corrected t_(83.85)_ = 4.208; p < 0.0001), AON (Welch-corrected t_(43.38)_ = 5.031; p < 0.0001), and aPIR (Welch-corrected t_(24.05)_ = 4.772; p < 0.0001) (**Fig. 2B**), whereas OXTR levels were decreased in the PFC (Welch-corrected t_(77.25)_ = 6.508; p < 0.0001), the AON (Welch-corrected t_(28.18)_ = 5.928; p < 0.0001) and the aPIR (t_(61)_ = 4.058; p < 0.001) (**Fig. 2C**). Our results indicate that the lack of MeCP2 causes alterations both in the IGF-1 and the OXT receptors in specific brain regions where KCC2 has been found to be altered. This suggests that the two systems could potentially be pharmacologically manipulated to ameliorate KCC2 imbalances in the RTT brain.

**Fig. 2.**
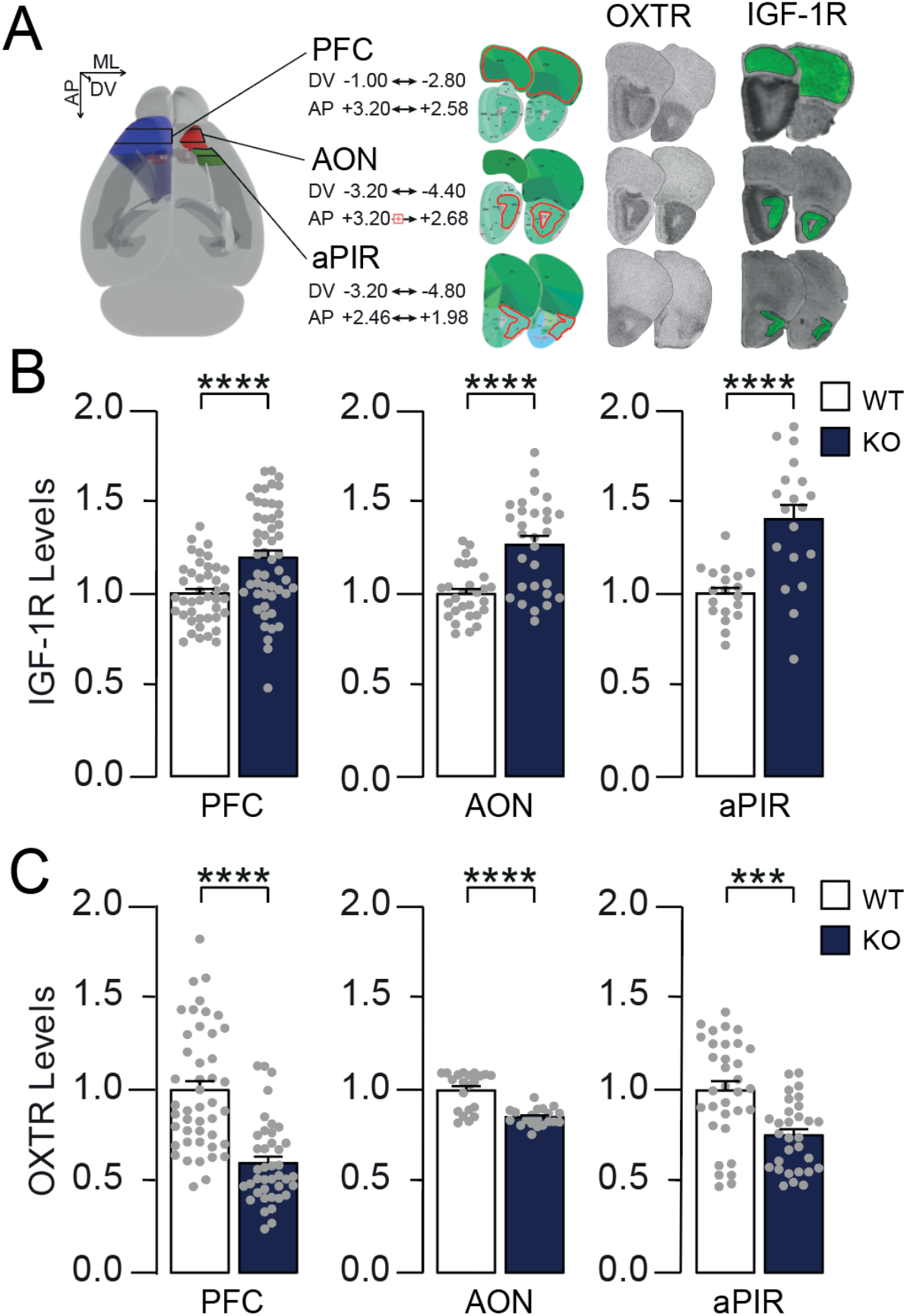
Brain areas with KCC2 reductions also display alterations in IGF-1R and OXTR expression. In PFC, AON and aPIR, KCC2 level reductions were accompanied by increased IGF-1R and decreased OXTR level. (**A**) Brain reconstruction showing the stereotactic coordinates of the coronal planes analyzed for each region. On the right, reference atlas coronal sections (Allen Brain Atlas) with matching representative infrared IGF-1R labelled and autoradiography labelled coronal brain sections. Areas outlined in red represent the regions of interest quantified in the analysis. (**B**) Bar graphs of IGF-1R levels (plotted as fold change over WT-Veh) displaying that IGF-1R was increased in the three KCC2-deficient areas in the MeCP2 KO mice; (**C**) Bar graphs of OXTR levels (plotted as fold change over WT-Veh) displaying that OXTR was consistently decreased in the three KCC2-deficient areas in the MeCP2 KO mice. Data expressed as mean ± SEM, single dots represent single observations from 3-4 animals per group, 2 or 3 coronal planes per brain region, 2-3 sections per coronal plane, left and right hemisphere data pooled together. ***p < 0.001, ****p < 0.0001 vs WT-Veh. Image credit for brain reference plates: Allen Institute.

### rhIGF-1 and OXT treatment rescue KCC2 expression in complementary brain regions

We next asked if rhIGF-1 or OXT treatment could rescue KCC2 levels in the key brain areas in symptomatic MeCP2 KO mice, and if this effect could be linked to the modulation of IGF-1R and OXTR expression levels. To address this, we injected mice (n = 3), via daily systemic intraperitoneal injections for 8-10 days, either with saline (control) or rhIGF-1 (2.5 mg/kg). In an additional set of mice (n = 3), we applied intranasal OXT (0.3 IU/administration, administered twice/day) (**Fig. 3A**). We followed doses shown to be effective for rescuing synaptic and behavioral phenotypes for rhIGF-1 application (Castro et al. 2014; Banerjee et al. 2016) or OXT treatment (Huang et al. 2014). When analyzing IGF-1R and OXTR receptor levels, we found a significant effect of treatment in all the three brain areas analyzed (one-way ANOVA, treatment effect on IGF-1R in the AON [F(3,84) = 16.09; p < 0.0001], aPIR [F(3,73) = 3.571; p < 0.001], PFC [F(3,147) = 4.657; p < 0.01]; one-way ANOVA treatment effect on OXTR in the AON [F(3,69) = 23.63; p < 0.0001], aPIR [F(3,103) = 7.752; p < 0.0001], and the PFC [F(3,135) = 18.32; p < 0.0001]). In particular, we found that rhIGF-1 rescued IGF-1R levels in the AON (p < 0.0001 versus MeCP2 KO-Vehicle; **Fig. 3B**) and OXTR levels in the PFC (p < 0.0001 versus MeCP2 KO-Vehicle; **Fig. 3C**). OXT treatment instead did not display any effects on IGF-1R and OXTR in any of the regions analyzed (**Fig. 3B** and **3C**). While infrared immunofluorescence analysis of KCC2 levels showed effects of treatment in all the three areas (one-way ANOVA treatment effect in the AON [F(3,76) = 21.45; p < 0.0001], aPIR [F(3,66) = 5.934; p < 0.01], and in the PFC [F(3,151) = 6.836; p < 0.001]), no modulation of KCC2 levels matching rhIGF-1’s action on the receptors was detected (**Fig. 3D**). More interestingly, however, rhIGF-1 and OXT displayed regional complementarity in rescuing KCC2. rhIGF-1 treatment normalized KCC2 expression in the aPIR (p < 0.05 versus MeCP2 KO-Vehicle; **Fig. 3D**), and OXT treatment rescued KCC2 expression in the AON (p < 0.0001 versus MeCP2 KO-Vehicle; **Fig. 3D**) and in the PFC (p < 0.001 versus MeCP2 KO-Vehicle; **Fig. 3D**). In the AON, where OXT effectively rescued KCC2 levels, rhIGF-1 treatment unexpectedly further decreased KCC2 expression (p<0.0001 versus WT-Vehicle; **Fig. 3D**). Overall, our analysis revealed that: 1) both IGF-1 and OXT receptors, in specific brain areas, can be pharmacologically modulated in MeCP2 KO mice, and in particular we found that OXTR is capable of being modulated by rhIGF-1 treatment in the PFC; 2) both rhIGF-1 or OXT treatments are effective in rescuing KCC2 levels in complementary brain regions, suggesting a potential combinatorial therapeutic strategy in MeCP2 KO mice (**Fig. 4**).

**Fig. 3.**
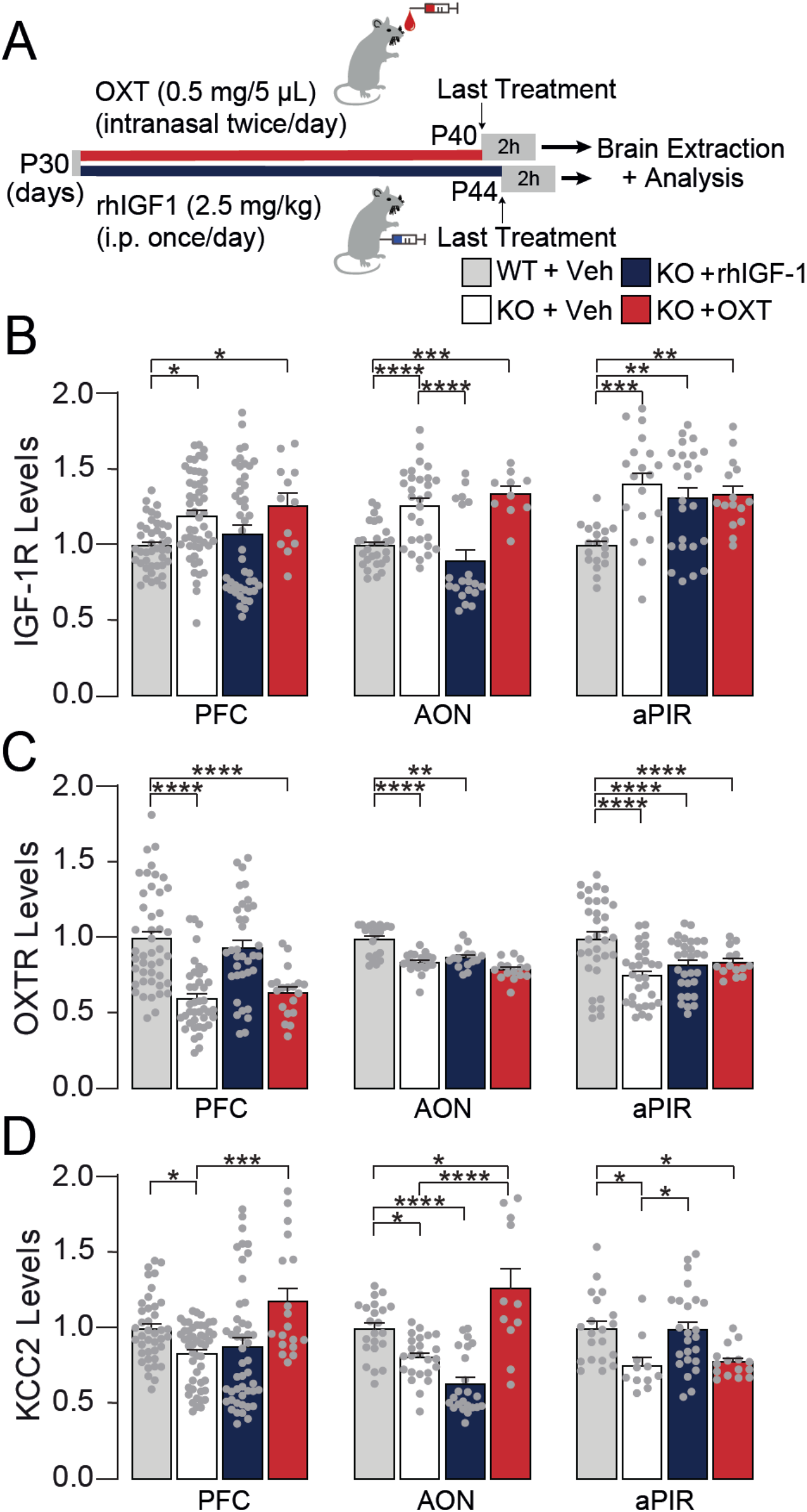
rhIGF-1 and OXT rescue KCC2 levels in complementary brain regions in MeCP2 KO mice. KCC2 levels are rescued independently by rhIGF-1 and OXT in MeCP2 KO animals, and these effects are not accompanied by rescue in the IGF-1R or OXTR levels, although rhIGF-1 is able to modulate both receptors in different areas. (**A**) A schematic diagram representing the treatment regime administered to the mice. (**B**) rhIGF-1 treatment normalized IGF-1R in the AON of MeCP2 KO mice, an effect not seen in PFC and aPIR, whereas OXT it did not modulate IGF-1R in any of the regions. (**C**) OXTR expression in the PFC is modulated by rhIGF-1 in MeCP2 KO mice, but no effect was seen in the AON or the aPIR. Moreover, OXTR levels were not modulated by OXT in any of the regions analyzed. (**D**) rhIGF-1 and OXT administration, instead, exerted different outcomes on KCC2 levels: indeed, rhIGF-1 was ineffective in the PFC, it diminished KCC2 in the AON, while it increased it up to WT-Veh levels in the aPIR. OXT treatment instead normalised KCC2 levels in the PFC and AON, but not in the aPIR. Data expressed as mean ± SEM, single dots represent single observations from 3-4 animals per group, 2 or 3 coronal planes per brain region, 2-3 sections per coronal plane, left and right hemisphere data pooled together. *p < 0.05, **p < 0.01, ***p < 0.001, ****p < 0.0001 vs WT-Veh or KO-Veh.

**Fig. 4.**
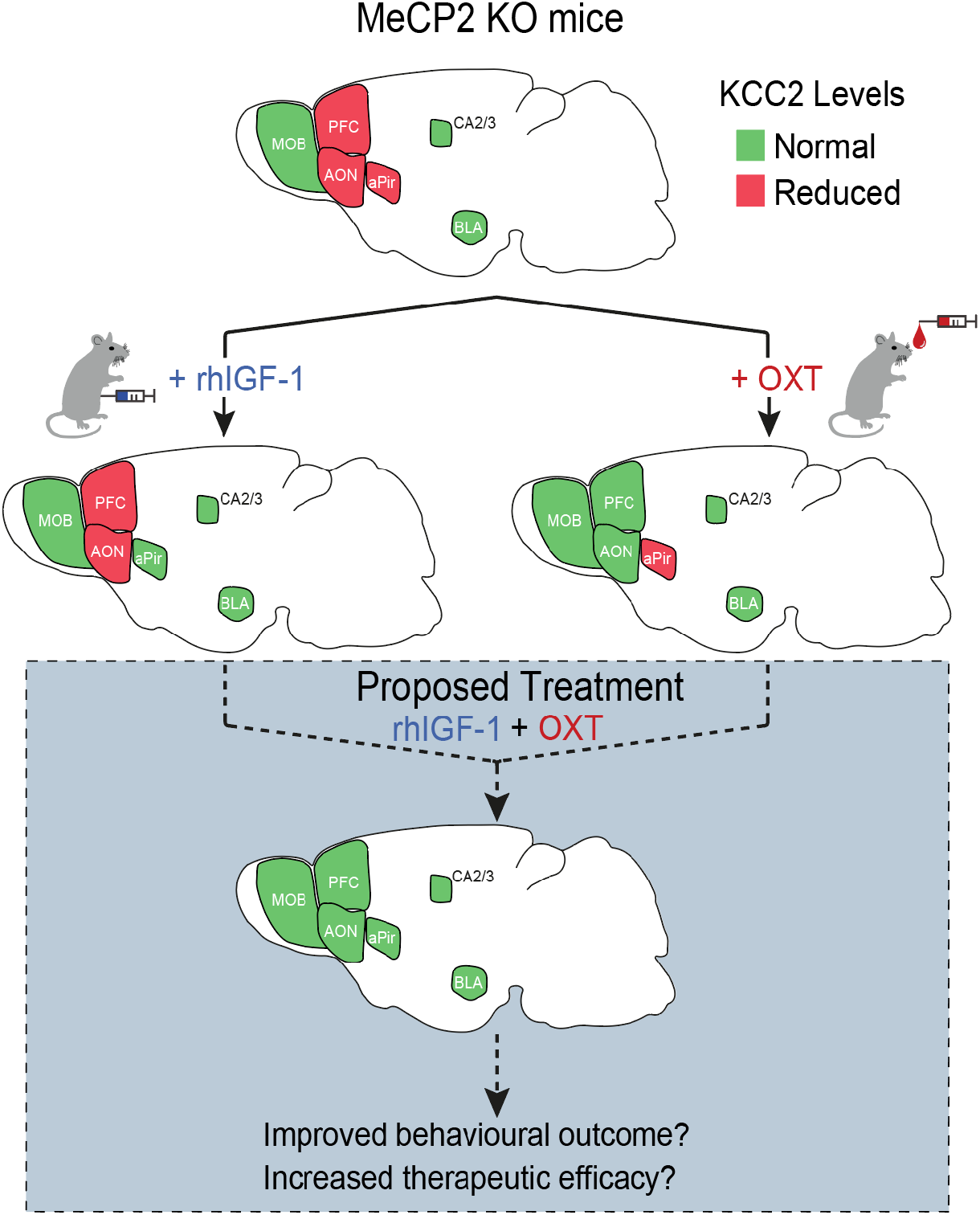
Potential for combined rhIGF-1 and OXT therapeutics in MeCP2 KO mice. Top: the lack of MeCP2 in mice induces regions-dependant deficits of KCC2. Areas in red display KCC2 reductions (AON, aPIR and PFC), whereas areas in green show normal levels of KCC2 (MOB, AONm, Hipp CA2/3, and BLA). Treatment with rhIGF-1 rescues KCC2 levels in the aPIR (Middle row, left), while treatment with OXT normalises KCC2 in the AON and PFC (Middle row, right). We therefore propose that a combined treatment of rhIGF-1 and OXT could lead to a more effective KCC2 normalization in the brain. If verified, this may represent a new, more efficacious treatment for RTT (Bottom row).

## Discussion

KCC2 is a transmembrane Cl^-^ transporter primarily involved in the maintenance of an appropriate GABAergic tone contributing to E/I balance. Dysregulation of E/I balance is a key feature of RTT pathophysiology, and is believed to delineate several of the RTT symptomatology including epilepsy, whose severity concurs to the clinical severity of the RTT phenotype (Glaze et al. 2010). In this study, we analyzed regional KCC2 expression levels in the MeCP2 KO mouse model of RTT. We show, for the first time, that KCC2 levels are significantly altered in multiple key brain areas. Such deficits are accompanied by alterations of the expression of IGF-1R and OXTR. Furthermore, we also show that rhIGF-1 treatment can rescue IGF-1R and OXTR levels in specific regions in symptomatic RTT mice. Finally, we demonstrate that rhIGF-1 and OXT exploit region-specific complementary rescue effects on KCC2 modulation in the brain of RTT mice.

### Circuit-specific KCC2 deficiency in MeCP2 KO mice

Throughout development, KCC2 transcription and expression is under the control of the MeCP2 protein. Interestingly, even though *Mecp2* mutations result in a global knockdown of the protein in male mice, KCC2 levels were selectively reduced in a region-specific manner reflecting circuit-specific dysfunction, as previously reported in c-Fos activity mapping studies (Kron et al. 2012). Even within the olfactory system, we found a reduction of KCC2 in the AON and aPIR, but not in other areas such as the MOB. Considering that KCC2 reductions are associated with higher seizure susceptibility, the specific KCC2-deficient brain areas could represent valid intervention points for the management of seizures in RTT patients. Our results are also consistent with the knowledge that KCC2 expression levels are regulated in a region-specific way during brain development (Wang et al. 2002).

In our study, we did not find significant reductions in KCC2 levels in the CA2/3 area of the hippocampus in MeCP2 KO male mice, differing with earlier observations reporting a reduction (El-Khoury et al. 2014). Interestingly, the study from El-Khoury and colleagues evaluated KCC2 levels using Western blot in the whole hippocampus, not in the isolated CA3. This suggests that in some brain areas KCC2 deficits could be confined to specific subregions, an aspect worth considering during preclinical study design. In this context, our results support the hypothesis that differences seen in KCC2 expression in some areas could be also developmental stage specific, an idea further corroborated by our finding of the absence of KCC2 deficits in MeCP2 KO P10 mice. KCC2 expression during development has been proved to be highly region and species-specific. In the mouse, Hsing and colleagues (Hsing et al. 2020) showed that in neocortical areas very low KCC2 expression can be detected at P0, whereas KCC2 signal in the piriform cortex can be detected as early as E15.5. Nonetheless both regions display similar levels of expression by P7. With this premises we quantified KCC2 in P10 mice, aiming at a developmental stage where not only the piriform cortex, but also prefrontal areas display detectable levels of KCC2. Indeed, we detected KCC2 in all our three regions of interest, i.e. the AON, the PFC and the aPIR. Considering that MeCP2 developmental expression is highly age and region-specific in itself, and that in the rat brain, MeCP2 expression increases dramatically in the first two postnatal weeks in the cortex and the hippocampus (Mullaney et al. 2004), it seemed reasonable that developmental differences in KCC2 levels could have been detectable by the age of P10. On the contrary, our KCC2 expression analysis did not display differences between WT and MeCP2 KO littermates in P10 mice. The idea that MeCP2 deficiency could impact preponderantly on KCC2 levels only at later stages of development is intriguing, especially in light of the delayed comparisons of RTT symptoms in mice and patients. This time and region-dependent expression of KCC2 could also make different brain regions and circuits vulnerable at different developmental stages. The mechanisms behind region-specific alteration of KCC2 in MeCP2 KO mice are unclear and require further investigation. This could be linked to either direct MeCP2 dysregulation effects on the KCC2 gene, or to indirect mechanisms involving, among others, OXT and IGF-1 signalling at the early stages of postnatal neurodevelopment, as discussed in the following sections.

### Altered OXT signalling in MeCP2 KO mice

To evaluate a possible contribution of a dysfunctional OXT signaling in RTT, we quantified the levels of the OXTR in the same brain areas where KCC2 level was significantly reduced. Our results show that, at least in the brain areas analyzed, OXTR is indeed dysregulated in MeCP2 KO mice. This finding is in line with reports of regional OXTR alterations in other murine models of neurodevelopment, such as the *Oprm1* KO, the valproic acid, the maternal immune activation, and the reelin heterozygous mouse models of autism (Liu et al. 2005; Gigliucci et al. 2014; Štefánik et al. 2015; Minakova et al. 2019). OXTR is modulated by several factors, the main being OXT itself. It is not known if there is a direct correlation between endogenous OXT production/release and OXTR expression levels, however chronic administration of OXT in adult mice can downregulate OXTR in several brain regions (Huang et al. 2014), raising the question whether an excessive production/release of endogenous OXT in MeCP2 KO mice may determine OXTR reductions. This hypothesis seems unlikely as recent reports suggest that the lack of MeCP2 does not affect cell density or distribution of OXT-positive neurons in the paraventricular and supraoptic nuclei of the hypothalamus, the two main production sites of OXT in the brain; moreover, MeCP2 deficiency seems to only marginally affect the distribution of OXT fibres in the brain (Martínez-Rodríguez et al. 2020). Nonetheless, a reduced level of OXTR could lead to impaired OXT signalling in specific brain regions, even in the absence of widespread alterations in hypothalamic OXT synthesis and local release. Another possibility for OXTR reductions in MeCP2 KO mice is that MeCP2 epigenetically or transcriptionally modulates some OXTR-modulating factors. For instance, it is known that OXTR is epigenetically modulated by Tet-1 (Towers et al. 2018), whose action on DNA can be inhibited by MeCP2 (Ludwig et al. 2017).

OXT administration at birth or in young adults displays therapeutic efficacy on social symptoms in mice models of neurodevelopmental pathologies, such as the *Oprm1* KO model of autism and the Magel2 KO model of Prader-Willy Syndrome (Gigliucci et al. 2014; Meziane et al. 2015; Bertoni et al. 2021). In our study, OXT treatment rescued KCC2 levels in two out of the three brain regions with KCC2 deficits, strongly suggesting a therapeutic potential of such an approach. In particular, OXT was active in the AON and in the PFC, in line with the well-established role of this neuropeptide as a master regulator of social behavior. The AON is a crucial feedback station in the olfactory system and it receives strong OXTergic innervation (Knobloch et al. 2012). It has recently been shown that OXT in this nucleus “sets the olfactory system into social receptiveness”, by enhancing the signal-to-noise ratio of odor responses, inducing olfactory exploration and by improving social recognition, and OXTR is necessary to exert these functions (Oettl et al. 2016). At the same time, the PFC is fundamental for discrimination of conspecifics’ affective state in mice (Scheggia et al. 2020), and OXTR in the PFC is pivotal for social recognition (Tan et al. 2019). It has recently been shown that OXTR activation in the nucleus accumbens is necessary for MDMA’s action to re-instate a window, otherwise closed, for social reward learning (Nardou et al. 2019). This suggests that OXT treatment could be a therapeutic strategy to ameliorate social deficits relevant to RTT symptomatology, similarly to what has been proposed for autistic children (Guastella et al. 2010; Auyeung et al. 2015). Another possibility is that OXT effects on KCC2 in these regions may be linked to the role of OXT on neuronal maturation.

In addition to differences in the level of expression of OXT receptors, other mechanisms can play a critical role in OXT’s regulation of KCC2 across brain regions. First, when binding OXTR, OXT recruits different intracellular G-protein coupled signaling pathways that can elicit opposite effects on neuronal activity, depending on the amount of peptide that is available extracellularly (Gravati et al., 2010; Eliava et al., 2016; Chini et al., 2017; Busnelli and Chini, 2018): at low OXT concentrations, the activated OXTR recruits the stimulatory Gq pathway, while higher concentrations of OXT induce the recruitment of the inhibitory G_i_ pathway, resulting in different final outcomes. Region-specific effects on OXT target(s) could thus be determined by the level of extracellular OXT, by the specific levels of G-protein isoforms and by cell-specific second messenger mediators, independently from the level of OXTR. Secondly, OXTR has been shown to be expressed in different neuronal cells as well as in astrocytes, potentially resulting in a variety of cellular outcomes. Finally, an intriguing hypothesis is that OXT could participate in regulating the release of IGF-1 itself, at least in some cells/regions, as shown in the periphery (Sirotkin et al., 2003), resulting in additional IGF-1 mediated effects on KCC2. Similarly, it has been shown that IGF-1 may inhibit supraoptic OXT neurons, possibly contributing to regulate OXT release within the brain (Ster et al. 2005).

### IGF-1 interaction with KCC2 in MeCP2 KO mice

When quantifying IGF-1R, we found that IGF-1R levels were consistently increased across the three brain regions analyzed. These increases could be secondary to reduced endogenous levels of circulating IGF-1 or IGF-1 binding protein (Schaevitz et al. 2010; Castro et al. 2014) or direct epigenetic repression of IGF-1R by MeCP2 during development. In MeCP2 mice, pharmacological treatment with rhIGF-1 normalized KCC2 levels region-selectively. In other studies based on exogenous IGF-1 administration or IGF-1R pharmacological inhibition, IGF-1 has been clearly implicated in the regulation of KCC2 expression and maturation of neurons in several brain areas including the hippocampus and the neocortex (Kelsch et al. 2001; Baroncelli et al. 2017). For example, Baroncelli et al. 2017 showed that blocking IGF-1 signalling by intracerebroventricular injections of the IGF-1 receptor antagonist JB1 prevented the effects of environmental enrichment linked to KCC2/NKCC1 function. However, to our knowledge, this is the first time that a region-selective effect of IGF-1 on KCC2 has been reported. In particular, the treatment was effective in the aPIR, which is a key region for olfactory discrimination and memory (Roesch et al. 2007). That rhIGF-1 treatment normalized KCC2 only in one brain area of the three analyzed was unexpected, however some speculations on the mechanisms behind it can be made. One possibility could be that sensory cortices are somehow more sensitive to rhIGF-1. Indeed, rhIGF-1 was shown to rescue KCC2 in MeCP2 KO mice in the visual cortex (Banerjee et al. 2016; Baroncelli et al. 2017). Sensory cortical functions are critical for neurodevelopment: stimuli from the environment activate and reinforce specific network connectivity and shape the animal’s perception of the world (Banerjee et al. 2021). IGF-1 is involved in experience-dependent plasticity in the visual cortex (Ciucci et al. 2007; Tropea et al. 2009; Scholl and Banerjee 2014), and the rescue of KCC2 in the aPIR suggests that a similar mechanism may occur in the piriform cortex. Interestingly, the aPIR is enriched in parvalbumin-expressing interneurons (Badowska-Szalewska et al. 2004), that express IGF-1R and are critical for RTT-linked reduced inhibition in the visual cortex of MeCP2 KO mice (Banerjee et al. 2016). Another possibility is that the lack of MeCP2 in sensory cortices makes them more immature, extending rhIGF-1’s therapeutic window, a prospect considered earlier (Banerjee et al. 2019). It has been shown that the loss of MeCP2 causes immaturity in selected areas of the olfactory system, including the piriform cortex (Martínez-Rodríguez et al., 2019). The KCC2 expression profile we report here is in line with this region-selective immaturity. Finally, the region-specific expression of IGF1R could be regulated by the local availability of IGF itself. It is interesting to note here that brain IGF-1 is predominantly produced peripherally, and that the serum IGF-1 can enter in selected by brain regions via an activity-dependent mechanism, suggesting the hypothesis that the effects of IGF-1 may also depend on the functional maturation of those areas (Aleman and Torres-Aleman 2009; Nishijima et al. 2010).

In conclusion, our data demonstrated that KCC2 levels are decreased in selective frontal brain regions in MeCP2 KO mice, and these are accompanied with similar alterations in OXTR and IGF-1R levels. OXT and rhIGF-1 displayed regional complementary effects on KCC2 expression: rhIGF-1 rescued KCC2 levels selectively in the aPIR, whereas OXT rescued levels in the AON and in the PFC. While the functional implications of this region-selectivity remain to be established, these findings strongly suggest that synergistic treatments of two important effectors may improve diverse aspects of the RTT symptomatology, and that a well-timed combination of these treatments may represent a powerful therapeutic strategy.

## Supporting information

Gigliucci_SupplMaterial_FINAL

## Acknowledgement

A.B. dedicates this work to the memory of his mentor Prof. Kshitish Majumdar. This work was supported by a H2020 Marie Skłodowska-Curie Actions fellowship (CIRCDYN, Grant number: 709288) and Brain and Behavior Research Foundation NARSAD Young Investigator award (Grant number 24941) to A.B.; Fritz Thyssen Foundation (Grant number 10.19.1.015MN) to B.C.; Brain and Behavior Research Foundation NARSAD Young Investigator award (Grant number 24947) to M.B.; Fondazione Umberto Veronesi fellowship (Grant number 3418) to V.G. We thank Drs. F. Helmchen and V. Broccoli for their support.

## Author contributions

A.B. and B.C. designed the study; V.G., A.B., J.T. and M.L. carried out experiments; M.W-S. conducted the gene network analysis; V.G., M.B. and B.C. analyzed and interpreted the molecular data; V.G. wrote the manuscript with B.C. and A.B. with comments from all authors. The authors declare no conflict of interest.

## Bibliography

Aleman A, Torres-Alemán I. 2009. Circulating insulin-like growth factor I and cognitive function: neuromodulation throughout the lifespan. Prog Neurobiol. 89(3):256–65.

Auyeung B, Lombardo M V., Heinrichs M, Chakrabarti B, Sule A, Deakin JB, Bethlehem RAI, Dickens L, Mooney N, Sipple JAN, Thiemann P, Baron-Cohen S. 2015. Oxytocin increases eye contact during a real-time, naturalistic social interaction in males with and without autism. Transl Psychiatry. 5(2):e507.

Badowska-Szalewska E, Klejbor I, Ludkiewicz B, Łuczyńska A, Domaradzka-Pytel B, Dziewiatkowski J, Moryś J. 2004. Parvalbumin containing neurons of the piriform cortex in open field stress - a developmental study in the rat. Folia Morphol (Warsz). 63(4):367–372.

Baker E, Stavropoulos KKM. 2020. The effects of oxytocin administration on individuals with ASD: Neuroimaging and behavioral evidence. Prog Mol Biol Transl Sci. 173:209–238.

Banerjee A, Castro J, Sur M. 2012. Rett syndrome: genes, synapses, circuits, and therapeutics. Front Psychiatry. 3:34.

Banerjee A, Miller MT, Li K, Sur M, Kaufmann WE. 2019. Towards a better diagnosis and treatment of Rett syndrome: a model synaptic disorder. Brain. 142(2):239–248.

Banerjee A, Parente G, Teutsch J, Lewis C, Voigt FF, Helmchen F. 2020. Value-guided remapping of sensory cortex by lateral orbitofrontal cortex. Nature. 585(7824):245–250.

Banerjee A, Rikhye R V, Breton-Provencher V, Tang X, Li C, Li K, Runyan CA, Fu Z, Jaenisch R, Sur M. 2016. Jointly reduced inhibition and excitation underlies circuit-wide changes in cortical processing in Rett syndrome. Proc Natl Acad Sci U S A. 113(46):E7287–E7296.

Banerjee A, Rikhye R V, Marblestone A. 2021. Reinforcement-guided learning in frontal neocortex: emerging computational concepts. Curr Opin Behav Sci. 38:133–140.

Baroncelli L, Cenni MC, Melani R, Deidda G, Landi S, Narducci R, Cancedda L, Maffei L, Berardi N. 2017. Early IGF-1 primes visual cortex maturation and accelerates developmental switch between NKCC1 and KCC2 Chloride transporters in enriched animals. Neuropharmacology. 113(Pt A):167–177.

Ben-Ari Y, Khalilov I, Kahle KT, Cherubini E. 2012. The GABA excitatory/inhibitory shift in brain maturation and neurological disorders. Neuroscientist. 18(5):467–486.

Bertoni A, Schaller F, Tyzio R, Gaillard S, Santini F, Xolin M, Diabira D, Vaidyanathan R, Matarazzo V, Medina I, Hammock E, Zhang J, Chini B, Gaiarsa JL, Muscatelli F 2021. Oxytocin administration in neonates shapes hippocampal circuitry and restores social behavior in a mouse model of autism. Mol Psychiatry. Online ahead of print.

Busnelli M, Chini B. 2018. Molecular Basis of Oxytocin Receptor Signalling in the Brain: What We Know and What We Need to Know. Curr Top Behav Neurosci. 35:3–29.

Castro J, Garcia RI, Kwok S, Banerjee A, Petravicz J, Woodson J, Mellios N, Tropea D, Sur M. 2014. Functional recovery with recombinant human IGF1 treatment in a mouse model of Rett Syndrome. Proc Natl Acad Sci U S A. 111(27):9941–9946.

Chini B, Verhage M, Grinevich V. The Action Radius of Oxytocin Release in the Mammalian CNS: From Single Vesicles to Behavior. 2017. Trends Pharmacol Sci. 38(11):982–991.

Ciucci F, Putignano E, Baroncelli L, Landi S, Berardi N, Maffei L. 2007. Insulin-like growth factor 1 (IGF-1) mediates the effects of enriched environment (EE) on visual cortical development. PLoS One. 2(5):e475.

Degano AL, Park MJ, Penati J, Li Q, Ronnett G V. 2014. MeCP2 is required for activity-dependent refinement of olfactory circuits. Mol Cell Neurosci. 59:63–75.

DeMayo MM, Song YJC, Hickie IB, Guastella AJ. 2017. A Review of the Safety, Efficacy and Mechanisms of Delivery of Nasal Oxytocin in Children: Therapeutic Potential for Autism and Prader-Willi Syndrome, and Recommendations for Future Research. Pediatr Drugs. 19(5):391–410.

Duarte ST, Armstrong J, Roche A, Ortez C, Pérez A, O’Callaghan M del M, Pereira A, Sanmartí F, Ormazábal A, Artuch R, Pineda M, García-Cazorla A, Pérez A, O’Callaghan Mdel M, Pereira A, Sanmartí F, Ormazábal A, Artuch R, Pineda M, García-Cazorla A. 2013. Abnormal expression of cerebrospinal fluid cation chloride cotransporters in patients with Rett syndrome. PLoS One. 8(7):e68851.

Eaton SL, Cumyn E, King D, Kline RA, Carpanini SM, Del-Pozo J, Barron R, Wishart TM. 2016. Quantitative imaging of tissue sections using infrared scanning technology. J Anat. 228(1):203–213.

Eliava M, Melchior M, Knobloch-Bollmann HS, Wahis J, da Silva Gouveia M, Tang Y, Ciobanu AC, Triana Del Rio R, Roth LC, Althammer F, Chavant V, Goumon Y, Gruber T, Petit-Demoulière N, Busnelli M, Chini B, Tan LL, Mitre M, Froemke RC, Chao MV, Giese G, Sprengel R, Kuner R, Poisbeau P, Seeburg PH, Stoop R, Charlet A, Grinevich V. 2016. A New Population of Parvocellular Oxytocin Neurons Controlling Magnocellular Neuron Activity and Inflammatory Pain Processing. Neuron. 89(6):1291–1304.

El-Khoury R, Panayotis N, Matagne V, Ghata A, Villard L, Roux JC. 2014. GABA and Glutamate Pathways Are Spatially and Developmentally Affected in the Brain of Mecp2-Deficient Mice. PLoS One. 9(3):e92169.

Feldman D, Banerjee A, Sur M. 2016. Developmental Dynamics of Rett Syndrome. Neural Plast. 2016:6154080.

Franklin, K. B. J.; Paxinos G. 2007. The mouse brain in stereotaxic coordinates - Third Ed.

Gigliucci V, Leonzino M, Busnelli M, Luchetti A, Palladino VS, D’Amato FR, Chini B. 2014. Region specific up-regulation of oxytocin receptors in the opioid Oprm1 (-/-) mouse model of autism. Front Pediatr. 2:91.

Glaze DG, Percy AK, Skinner S, Motil KJ, Neul JL, Barrish JO, Lane JB, Geerts SP, Annese F, Graham J, McNair L, Lee HS. 2010. Epilepsy and the natural history of Rett syndrome. Neurology. 74(11):909–912.

Gravati M, Busnelli M, Bulgheroni E, Reversi A, Spaiardi P, Parenti M, Toselli M, Chini B. 2010. Dual modulation of inward rectifier potassium currents in olfactory neuronal cells by promiscuous G protein coupling of the oxytocin receptor. J Neurochem. 114(5):1424–35.

Guastella AJ, Einfeld SL, Gray KM, Rinehart NJ, Tonge BJ, Lambert TJ, Hickie IB. 2010. Intranasal Oxytocin Improves Emotion Recognition for Youth with Autism Spectrum Disorders. Biol Psychiatry. 67(7):692–694.

Guy J, Hendrich B, Holmes M, Martin JE, Bird A. A mouse Mecp2-null mutation causes neurological symptoms that mimic Rett syndrome. Nat Genet. 2001 Mar;27(3):322–6.

Hinz L, Torrella Barrufet J, Heine VM. 2019. KCC2 expression levels are reduced in post mortem brain tissue of Rett syndrome patients. Acta Neuropathol Commun. 7(1):196.

Howell CJ, Sceniak MP, Lang M, Krakowiecki W, Abouelsoud FE, Lad SU, Yu H, Katz DM. 2017. Activation of the medial prefrontal cortex reverses cognitive and respiratory symptoms in a mouse model of Rett syndrome. eNeuro. 4(6):ENEURO.0277-17.2017.

Hsing HW, Zhuang ZH, Niou ZX, Chou SJ. 2020. Temporal Differences in Interneuron Invasion of Neocortex and Piriform Cortex during Mouse Cortical Development. Cereb Cortex. 30(5):3015–3029.

Hsu WL, Ma YL, Liu YC, Tai DJC, Lee EHY. 2020. Restoring Wnt6 signaling ameliorates behavioral deficits in MeCP2 T158A mouse model of Rett syndrome. Sci Rep. 10(1):1074.

Huang H, Michetti C, Busnelli M, Managò F, Sannino S, Scheggia D, Giancardo L, Sona D, Murino V, Chini B, Scattoni ML, Papaleo F. 2014. Chronic and acute intranasal oxytocin produce divergent social effects in mice. Neuropsychopharmacology. 39(5):1102–1114.

Kelsch W, Hormuzdi S, Straube E, Lewen A, Monyer H, Misgeld U. 2001. Insulin-like growth factor 1 and a cytosolic tyrosine kinase activate chloride outward transport during maturation of hippocampal neurons. J Neurosci. 21(21):8339–8347.

Khwaja OS, Ho E, Barnes K V., O’Leary HM, Pereira LM, Finkelstein Y, Nelson CA, Vogel-Farley V, DeGregorio G, Holm IA, Khatwa U, Kapur K, Alexander ME, Finnegan DM, Cantwell NG, Walco AC, Rappaport L, Gregas M, Fichorova RN, Shannon MW, Sur M, Kaufmann WE. 2014. Safety, pharmacokinetics, and preliminary assessment of efficacy of mecasermin (recombinant human IGF-1) for the treatment of Rett syndrome. Proc Natl Acad Sci U S A. 111(12):4596–4601.

Knobloch HS, Charlet A, Hoffmann LC, Eliava M, Khrulev S, Cetin AH, Osten P, Schwarz MK, Seeburg PH, Stoop R, Grinevich V. 2012. Evoked axonal oxytocin release in the central amygdala attenuates fear response. Neuron. 73(3):553–566.

Kron M, Howell CJ, Adams IT, Ransbottom M, Christian D, Ogier M, Katz DM. 2012. Brain activity mapping in Mecp2 mutant mice reveals functional deficits in forebrain circuits, including key nodes in the default mode network, that are reversed with ketamine treatment. J Neurosci. 32(40):13860–13872.

Krupp DR, Barnard RA, Duffourd Y, Evans SA, Mulqueen RM, Bernier R, Rivière JB, Fombonne E, O’Roak BJ. 2017. Exonic Mosaic Mutations Contribute Risk for Autism Spectrum Disorder. Am J Hum Genet. 101(3):369–390.

Leonzino M, Busnelli M, Antonucci F, Verderio C, Mazzanti M, Chini B. 2016. The Timing of the Excitatory-to-Inhibitory GABA Switch Is Regulated by the Oxytocin Receptor via KCC2. Cell Rep. 15(1):96–103.

Li Y, Shen M, Stockton ME, Zhao X. 2019. Hippocampal deficits in neurodevelopmental disorders. Neurobiol Learn Mem. 165:106945.

Liu W, Pappas GD, Carter CS. 2005. Oxytocin receptors in brain cortical regions are reduced in haploinsufficient (+/-) reeler mice. Neurol Res. 27(4):339–345.

Ludwig AK, Zhang P, Hastert FD, Meyer S, Rausch C, Herce HD, Müller U, Lehmkuhl A, Hellmann I, Trummer C, Storm C, Leonhardt H, Cardoso MC. 2017. Binding of MBD proteins to DNA blocks Tet1 function thereby modulating transcriptional noise. Nucleic Acids Res. 45(5):2438–2457.

Martínez-Rodríguez E, Martín-Sánchez A, Coviello S, Foiani C, Kul E, Stork O, Martínez-García F, Nacher J, Lanuza E, Santos M, Agustín-Pavón C. 2019. Lack of MeCP2 leads to region-specific increase of doublecortin in the olfactory system. Brain Struct Funct. 224(4):1647–1658.

Martínez-Rodríguez E, Martín-Sánchez A, Kul E, Bose A, Martínez-Martínez FJ, Stork O, Martínez-García F, Lanuza E, Santos M, Agustín-Pavón C. 2020. Male-specific features are reduced in Mecp2-null mice: analyses of vasopressinergic innervation, pheromone production and social behaviour. Brain Struct Funct. 225(7):2219–2238.

Meziane H, Schaller F, Bauer S, Villard C, Matarazzo V, Riet F, Guillon G, Lafitte D, Desarmenien MG, Tauber M, Muscatelli F. 2015. An Early Postnatal Oxytocin Treatment Prevents Social and Learning Deficits in Adult Mice Deficient for Magel2, a Gene Involved in Prader-Willi Syndrome and Autism. Biol Psychiatry. 78(2):85–94.

Minakova E, Lang J, Medel-Matus JS, Gould GG, Reynolds A, Shin D, Mazarati A, Sankar R. 2019. Melanotan-II reverses autistic features in a maternal immune activation mouse model of autism. PLoS One. 14(1):e0210389.

Mullaney BC, Johnston MV, Blue ME. 2004. Developmental expression of methyl-CpG binding protein 2 is dynamically regulated in the rodent brain. Neuroscience. 123(4):939–49.

Nardou R, Lewis EM, Rothhaas R, Xu R, Yang A, Boyden E, Dölen G. 2019. Oxytocin-dependent reopening of a social reward learning critical period with MDMA. Nature. 569(7754):116–120.

Nishijima T, Piriz J, Duflot S, Fernandez AM, Gaitan G, Gomez-Pinedo U, Verdugo JM, Leroy F, Soya H, Nuñez A, Torres-Aleman I. 2010. Neuronal activity drives localized blood-brain-barrier transport of serum insulin-like growth factor-I into the CNS. Neuron. 67(5):834–46.

O’Leary HM, Kaufmann WE, Barnes K V., Rakesh K, Kapur K, Tarquinio DC, Cantwell NG, Roche KJ, Rose SA, Walco AC, Bruck NM, Bazin GA, Holm IA, Alexander ME, Swanson LC, Baczewski LM, Mayor Torres JM, Nelson CA, Sahin M. 2018. Placebo-controlled crossover assessment of mecasermin for the treatment of Rett syndrome. Ann Clin Transl Neurol. 5(3):323–332.

Oettl LL, Ravi N, Schneider M, Scheller MF, Schneider P, Mitre M, da Silva Gouveia M, Froemke RC, Chao M V., Young WS, Meyer-Lindenberg A, Grinevich V, Shusterman R, Kelsch W. 2016. Oxytocin Enhances Social Recognition by Modulating Cortical Control of Early Olfactory Processing. Neuron. 90(3):609–621.

Pini G, Congiu L, Benincasa A, DiMarco P, Bigoni S, Dyer AH, Mortimer N, Della-Chiesa A, O’Leary S, McNamara R, Mitchell KJ, Gill M, Tropea D. 2016. Illness Severity, Social and Cognitive Ability, and EEG Analysis of Ten Patients with Rett Syndrome Treated with Mecasermin (Recombinant Human IGF-1). Autism Res Treat. 2016:5073078.

Pozzi D and Chini B 2019 Quest for pharmacological regulators of KCC2 In: Neuronal Chloride Transporters in Health and Disease, (Ed: Xin Tan), Academic Press (pp.709–727)

Roesch MR, Stalnaker TA, Schoenbaum G. 2007. Associative encoding in anterior piriform cortex versus orbitofrontal cortex during odor discrimination and reversal learning. Cereb Cortex. 17(3):643–652.

Samaco RC, McGraw CM, Ward CS, Sun Y, Neul JL, Zoghbi HY. Female Mecp2(+/-) mice display robust behavioral deficits on two different genetic backgrounds providing a framework for pre-clinical studies. Hum Mol Genet. 2013 Jan 1;22(1):96–109.

Sceniak MP, Lang M, Enomoto AC, James Howell C, Hermes DJ, Katz DM. 2016. Mechanisms of Functional Hypoconnectivity in the Medial Prefrontal Cortex of Mecp2 Null Mice. Cereb Cortex. 26(5):1938–1956.

Schaevitz LR, Moriuchi JM, Nag N, Mellot TJ, Berger-Sweeney J. 2010. Cognitive and social functions and growth factors in a mouse model of Rett syndrome. Physiol Behav. 100(3):255–263.

Scheggia D, Managò F, Maltese F, Bruni S, Nigro M, Dautan D, Latuske P, Contarini G, Gomez-Gonzalo M, Requie LM, Ferretti V, Castellani G, Mauro D, Bonavia A, Carmignoto G, Yizhar O, Papaleo F. 2020. Somatostatin interneurons in the prefrontal cortex control affective state discrimination in mice. Nat Neurosci. 23(1):47–60.

Scholl B, Banerjee A. 2014. Synaptic correlates of binocular convergence: Just a coincidence? J Neurosci. 34(27):8931–8933.

Sirotkin AV, Florkovicova I, Makarevich AV, Schaeffer HJ, Kotwica J, Marnet PG, Sanislo P. 2003. Oxytocin mediates some effects of insulin-like growth factor-I on porcine ovarian follicles. J Reprod Dev. 49(2):141–9.

Štefánik P, Olexová L, Kršková L. 2015. Increased sociability and gene expression of oxytocin and its receptor in the brains of rats affected prenatally by valproic acid. Pharmacol Biochem Behav. 131:42–50.

Stödberg T, McTague A, Ruiz AJ, Hirata H, Zhen J, Long P, Farabella I, Meyer E, Kawahara A, Vassallo G, Stivaros SM, Bjursell MK, Stranneheim H, Tigerschiöld S, Persson B, Bangash I, Das K, Hughes D, Lesko N, Lundeberg J, Scott RC, Poduri A, Scheffer IE, Smith H, Gissen P, Schorge S, Reith MEA, Topf M, Kullmann DM, Harvey RJ, Wedell A, Kurian MA. 2015. Mutations in SLC12A5 in epilepsy of infancy with migrating focal seizures. Nat Commun. 6:8038.

Tan Y, Singhal SM, Harden SW, Cahill KM, Nguyen DTM, Colon-Perez LM, Sahagian TJ, Thinschmidt JS, De Kloet AD, Febo M, Frazier CJ, Krause EG. 2019. Oxytocin receptors are expressed by glutamatergic prefrontal cortical neurons that selectively modulate social recognition. J Neurosci. 39(17):3249–3263.

Tang X, Drotar J, Li K, Clairmont CD, Brumm AS, Sullins AJ, Wu H, Liu XS, Wang J, Gray NS, Sur M, Jaenisch R. 2019. Pharmacological enhancement of KCC2 gene expression exerts therapeutic effects on human Rett syndrome neurons and Mecp2 mutant mice. Sci Transl Med. 11(503) eaau0164.

Tang X, Kim J, Zhou L, Wengert E, Zhang L, Wu Z, Carromeu C, Muotri AR, Marchetto MCN, Gage FH, Chen G. 2016. KCC2 rescues functional deficits in human neurons derived from patients with Rett syndrome. Proc Natl Acad Sci U S A. 113(3):751–756.

Towers AJ, Tremblay MW, Chung L, Li XL, Bey AL, Zhang W, Cao X, Wang X, Wang P, Duffney LJ, Siecinski SK, Xu S, Kim Y, Kong X, Gregory S, Xie W, Jiang YH. 2018. Epigenetic dysregulation of Oxtr in Tet1-deficient mice has implications for neuropsychiatric disorders. JCI insight. 3(23):e120592.

Tropea D, Giacometti E, Wilson NR, Beard C, McCurry C, Fu DD, Flannery R, Jaenisch R, Sur M. 2009. Partial reversal of Rett Syndrome-like symptoms in MeCP2 mutant mice. Proc Natl Acad Sci U S A. 106(6):2029–2034.

Tyzio R, Nardou R, Ferrari DC, Tsintsadze T, Shahrokhi A, Eftekhari S, Khalilov I, Tsintsadze V, Brouchoud C, Chazal G, Lemonnier E, Lozovaya N, Burnashev N, Ben-Ari Y. 2014. Oxytocin-mediated GABA inhibition during delivery attenuates autism pathogenesis in rodent offspring. Science. 343(6171):675–679.

Wang C, Shimizu-Okabe C, Watanabe K, Okabe A, Matsuzaki H, Ogawa T, Mori N, Fukuda A, Sato K. 2002. Developmental changes in KCC1, KCC2, and NKCC1 mRNA expressions in the rat brain. Dev Brain Res. 139(1):59–66.

