## Supplementary material for "Region-specific KCC2 rescue by rhIGF-1 and oxytocin in a mouse model of Rett syndrome": Gigliucci_SupplMaterial_FINAL

Valentina Gigliucci[1], Jasper Teutsch[2,3], Marc Woodbury-Smith[2], Mirko Luoni[4], Marta Busnelli[1,5], Bice Chini[1,5†], and Abhishek Banerjee[2,3†]

[1]CNR Institute of Neuroscience, Milan, Italy
[2]Neuroscience Theme, Biosciences Institute, Newcastle University, United Kingdom
[3]University of Zurich, Zurich, Switzerland
[4]San Raffaele Scientific Institute, Milan, Italy
[5]NeuroMi Milan Center for Neuroscience, Milan, Italy

**†Correspondence:**

Abhishek Banerjee

and

Bice Chini

### Supplementary Methods

#### Gene co-expression network analysis.

To characterise the co-expression network for *SLC12A5*, we followed the steps shown in **Supplementary Fig. 1A.** We used RNAseq gene expression data from the BrainSpan atlas of the developing human brain (Hawrylycz et al. 2012), and our analyses were conducted on brain samples from subjects between the $6^{th}$ post-conception week and 8 years of age. We excluded older samples to ensure our networks captured the early developing brain as Rett syndrome and the other neurodevelopmental disorders we are interested in all have onset in these early years. After excluding samples with RNA Integrity Number (RIN) less than 7.5, gene-level reads per kilobase million (RPKM) were normalised for GC content using procedures implemented in the R cqn package (Hansen et al. 2012), and all genes with RPKM less than 1 in 50% or more of samples were removed, i.e. those genes that were unexpressed across more than 50% of samples. The RPKM expression matrix was then loaded into R for co-expression analysis using WGCNA (Langfelder and Horvath 2008). Co-expression similarity between pairs of genes is represented by an adjacency matrix, with adjacency scores calculated from correlation between normalized gene expression with these scores then transformed according to the soft-thresholding procedure described in the WGCNA protocol. Hierarchical clustering was then used to define clusters of strongly interconnected genes. A minimum clustering threshold was set at 30 genes per cluster, and the Dynamic Tree Cut method was used to set dendogram height and thereby modules of closely expressed genes were generated. We then extracted the module containing *SLC12A5* to examine its characteristics as discussed subsequently.

We were interested in whether the co-expression network for KCC2 (*SLC12A5*) was enriched for neurodevelopmental genes, and so therefore performed an over-representation analyses using the R GeneOverlap package. This performs a hypergeometric test (i.e., a Fisher's exact test) to calculate the p-value in relation to the intersection between two gene lists. The ‘gene universe’ was set as all those genes included in the WGCNA analysis. Lists of ASD, ID and epilepsy implicated genes were derived from DisGenet (www.disgenet.org), a regularly updated collection of genes and variants associated with human diseases (Piñero et al. 2020). Additionally, we identified genes from SFARI that were classified Category 1 and 2 (high confidence and strong candidate sets). For each set of these genes we investigated whether a greater proportion were in the *SLC12A5* network than by chance. In order to further expound the functional role of *SLC12A5* and its interacting partners, we also investigated functional enrichment (over-representation) in the *SLC12A5* module for other gene sets (including GO pathways, KEGG, Reactome, and WikiPathways) using g:profiler (Raudvere et al. 2019).

### Supplementary Data

**Wild-type and MeCP2 KO mice do not show left/right hemispheres differences in regional KCC2, IGF-1R and OXTR expression.** Levels of KCC2, IGF-1R and OXTR from left and right hemispheres for each brain area and each experimental group were compared by Student's *t*-test analysis. No significant differences were highlighted by these analyses, as reported in **Supplementary tables 1 and 2**.

**Supplementary Table 1:** KCC2 expression does not show left/right hemisphere differences in any of the brain regions analysed (Multiple *t*-test after Sidak-Bonferroni correction, alpha = 5.0%).

| | WT-Veh | | | KO-Veh | | |
|---|---|---|---|---|---|---|
| **Brain area** | Right hemisphere (mean ± SEM) | Left hemisphere (mean ± SEM) | *p* value | Right hemisphere (mean ± SEM) | Left hemisphere (mean ± SEM) | *p* value |
| MOB | 115.4 ± 8.118 | 118.7 ± 7.451 | 0.7679, ns | 120.7 ± 5.817 | 120.7 ± 9.948 | 0.9950, ns |
| AON | 185.5 ± 9.999 | 188.2 ± 11.39 | 0.8674, ns | 241.1 ± 48.54 | 150.5 ± 8.227 | 0.0882, ns |
| AONm | 117.0 ± 11.52 | 130.5 ± 12.29 | 0.4341, ns | 141.6 ± 16.29 | 130.0 ± 15.04 | 0.6103, ns |
| aPIR | 169.5 ± 12.21 | 141.0 ± 7.316 | 0.0749, ns | 140.0 ± 19.27 | 109.9 ± 3.179 | 0.2250, ns |
| PFC | 236.1 ± 11.83 | 245.5 ± 13.98 | 0.6076, ns | 257.6 ± 28.56 | 271.4 ± 33.19 | 0.7527, ns |
| BLA | 84.09 ± 17.41 | 114.7 ± 25.60 | 0.3469, ns | 95.59 ± 24.26 | 88.49 ± 16.64 | 0.8124, ns |
| Hipp CA2/3 | 108.8 ± 17.92 | 116.0 ± 18.40 | 0.7828, ns | 88.33 ± 13.06 | 78.16 ± 16.11 | 0.6246, ns |

**Supplementary Table 2:** KCC2, IGF-1R and OXTR levels do not show left/right hemisphere differences in any of the brain regions analyzed (Multiple *t*-test after Sidak-Bonferroni correction, alpha = 5.0%, each target analyzed separately).

**KCC2 (Signal/pixel)**

| | PFC | | | AON | | | aPIR | | |
|---|---|---|---|---|---|---|---|---|---|
| Group | Right hemisphere (mean ± SEM) | Left hemisphere (mean ± SEM) | *p* value | Right hemisphere (mean ± SEM) | Left hemisphere (mean ± SEM) | *p* value | Right hemisphere (mean ± SEM) | Left hemisphere (mean ± SEM) | *p* value |
| WT-Veh | 236.1 ± 11.83 | 245.5 ± 13.98 | 0.6076, ns | 185.5 ± 9.999 | 188.2 ± 11.39 | 0.8674, ns | 169.5 ± 12.21 | 141.0 ± 7.316 | 0.0749, ns |
| KO-Veh | 257.6 ± 28.56 | 271.4 ± 33.19 | 0.7527, ns | 241.1 ± 48.54 | 150.5 ± 8.227 | 0.0882, ns | 140.0 ± 19.27 | 109.9 ± 3.179 | 0.2250, ns |
| KO-rhIGF-1 | 213.5 ± 19.91 | 217.4 ± 20.63 | 0.8936, ns | 116.5 ± 12.54 | 120.3 ± 10.48 | 0.8174, ns | 142.7 ± 10.44 | 171.7 ± 13.10 | 0.0935, ns |
| KO-OXT | 281.9 ± 29.50 | 297.5 ± 34.72 | 0.7358, ns | 221.9 ± 41.28 | 255.1 ± 27.18 | 0.5385, ns | 124.7 ± 6.747 | 119.6 ± 4.965 | 0.5528, ns |

**IGF-1R (Signal/pixel)**

| | PFC | | | AON | | | aPIR | | |
|---|---|---|---|---|---|---|---|---|---|
| Group | Right hemisphere (mean ± SEM) | Left hemisphere (mean ± SEM) | *p* value | Right hemisphere (mean ± SEM) | Left hemisphere (mean ± SEM) | *p* value | Right hemisphere (mean ± SEM) | Left hemisphere (mean ± SEM) | *p* value |
| WT-Veh | 127.8 ± 4.251 | 131.0 ± 5.082 | 0.6340, ns | 115.3 ± 3.577 | 122.0 ± 5.263 | 0.2844, ns | 88.57 ± 3.410 | 92.50 ± 5.428 | 0.5279, ns |
| KO-Veh | 152.3 ± 6.294 | 159.0 ± 8.766 | 0.5239, ns | 148.4 ± 7.534 | 151.2 ± 8.313 | 0.8044, ns | 119.0 ± 8.225 | 116.0 ± 22.42 | 0.8802, ns |
| KO-rhIGF-1 | 138.0 ± 10.36 | 141.1 ± 11.74 | 0.8460, ns | 104.5 ± 9.819 | 108.4 ± 14.56 | 0.8228, ns | 116.4 ± 8.459 | 121.2 ± 10.10 | 0.7136, ns |
| KO-OXT | 163.1 ± 9.894 | 160.2 ± 12.57 | 0.8601, ns | 156.4 ± 8.813 | 162.9 ± 6.204 | 0.6042, ns | 126.6 ± 6.787 | 108.7 ± 5.371 | 0.1092, ns |

**OXTR (nCi/mg tissue equivalent)**

| | PFC | | | AON | | | aPIR | | |
|---|---|---|---|---|---|---|---|---|---|
| Group | Right hemisphere (mean ± SEM) | Left hemisphere (mean ± SEM) | *p* value | Right hemisphere (mean ± SEM) | Left hemisphere (mean ± SEM) | *p* value | Right hemisphere (mean ± SEM) | Left hemisphere (mean ± SEM) | *p* value |
| WT-Veh | 0.1758 ± 0.011 | 0.1830 ± 0.017 | 0.7182, ns | 0.9576 ± 0.046 | 0.9471 ± 0.040 | 0.8664, ns | 0.5866 ± 0.051 | 0.6275 ± 0.052 | 0.5780, ns |
| KO-Veh | 0.1112 ± 0.011 | 0.09871 ± 0.008 | 0.3425, ns | 0.7405 ± 0.010 | 0.7862 ± 0.023 | 0.0955, ns | 0.4265 ± 0.034 | 0.4241 ± 0.032 | 0.9594, ns |
| KO-rhIGF-1 | 0.1462 ± 0.015 | 0.1853 ± 0.014 | 0.0690, ns | 0.8007 ± 0.027 | 0.7735 ± 0.035 | 0.7483, ns | 0.4654 ± 0.031 | 0.4802 ± 0.032 | 0.7404, ns |
| KO-OXT | 0.1115 ± 0.010 | 0.1113 ± 0.009 | 0.9862, ns | 0.6926 ± 0.031 | 0.6957 ± 0.024 | 0.9397, ns | 0.5144 ± 0.033 | 0.4575 ± 0.015 | 0.1115, ns |

**Gene co-expression network analysis.** We investigated the co-expression properties of the *SLC12A5* co-expression module in the developing human PFC, to examine enrichment for gene ontology terms and genes implicated in the three neurodevelopmental disorders most strongly associated with RTT, i.e., epilepsy, ASD, and ID (**Supplementary Fig. 1A-B**). Using expression data from the developing human brain from BrainSpan (Hawrylycz et al. 2012), we found that the co-expression network of KCC2 comprised 424 interacting partners (**Supplementary Table 3**). Confirming our hypothesis, a number of key neurodevelopmental genes do appear in this network, including *NRXN1*, *SHANK1*, *DLGAP1*, *RBFOX3* and others. We investigated gene set over-representation for the 424 genes in *SLC12A5*'s network with ASD, ID and epilepsy gene sets curated from different sources as discussed in the Supplementary Methods. At first, we examined SFARI Category 1 and 2 (high confidence and strong candidate sets) ASD genes (Abrahams et al. 2013), but significance was not demonstrated. However, when analyzing DisGenet curated gene sets (Piñero et al. 2020) for ASD, ID and epilepsy there were significant overlaps with the *SLC12A5* network ($p = 3.7e-03$, $p = 9.9e-03$ and $p = 4.9e-15$ for ASD, ID and epilepsy respectively), supporting the hypothesis that *SLC12A5* and its interacting partners have a key role in these neurodevelopmental disorders (**Supplementary Fig. 1B**).

Next, we extended our analysis by undertaking gene set over-representation using g:profiler (Raudvere et al. 2019) for gene ontology derived curated gene sets, with Bonferroni correction for multiple testing. As we expected, significant over-representation was observed for key synaptic gene sets including GO:CC synaptic membrane protein genes ($p = 5.8$ x10e-18), postsynaptic density protein genes ($p = 2.4 \times 10e-14$) (**Supplementary Fig. 3**), and GO:BP learning and memory ($p = 1.7 \times 10e-11$) (**Supplementary Fig. 4**). In line with its transporter function, other over-represented gene sets included ion transmembrane transporter activity and cell-cell signalling molecules (**Supplementary Fig. 5**). Taken together, these results highlight that *SLC12A5*'s interacting partners are largely synaptic and transporter proteins, consistent with their predicted role in disorders such as ASD and epilepsy. Gene Ontology (GO) enrichment analysis was performed for SLC12A5 and its co-expression network, with significant GO categories, is provided as the supplementary Excel file (**Supplementary Table 3**).

### Supplementary Figures

#### Supplementary Fig. 1

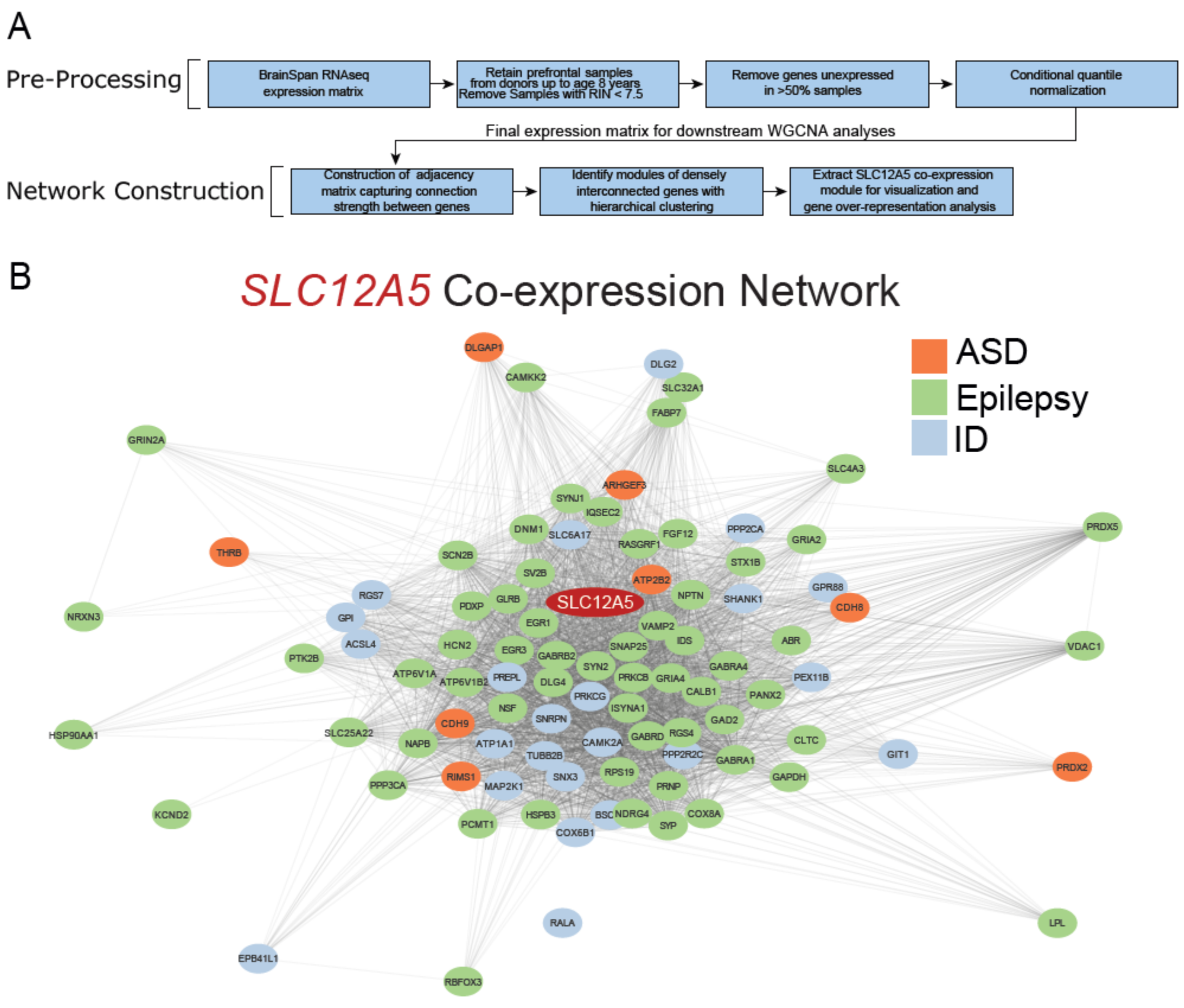


**Supplementary Fig. 1. *SLC12A5* co-expression Network in the developing human PFC. (A)** Schematic representation of the gene co-expression network analysis workflow; **(B)** The gene co-expression network for *SLC12A5* (red oval). The cloud of representative genes highlighted in the figure shows that the network is enriched in genes linked to three neurodevelopmental disorders most strongly associated with RTT: ASD (orange), epilepsy (green), and ID (light blue).

### Supplementary Fig. 2

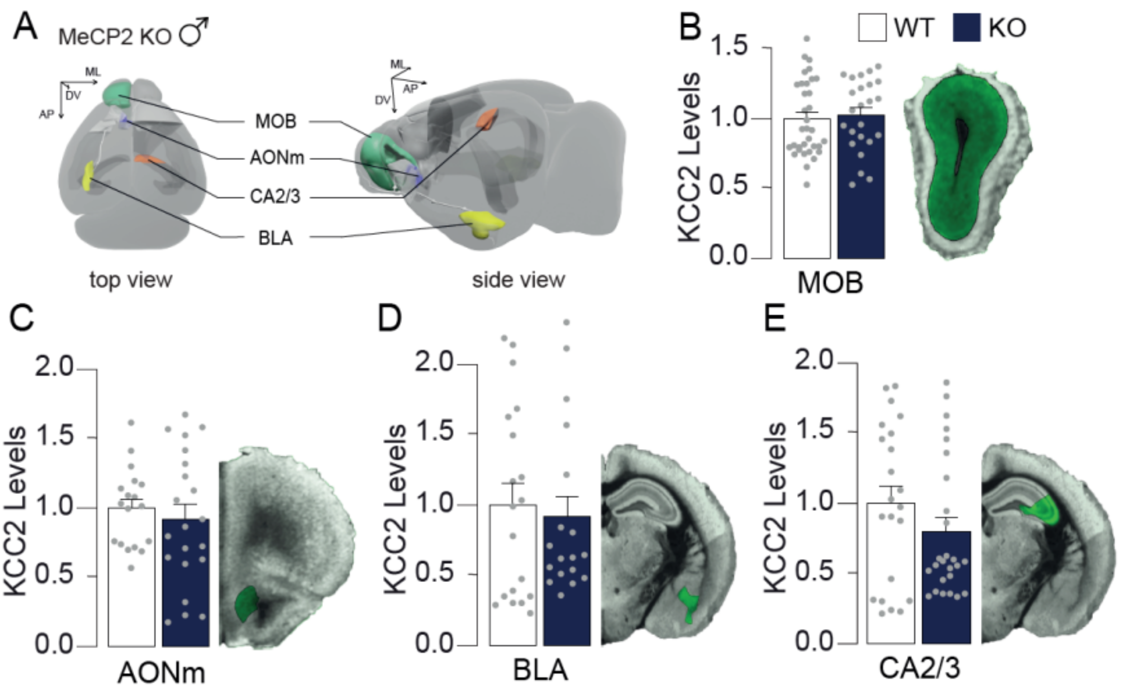


**Supplementary Fig. 2. Several brain regions in P44 male MeCP2 KO mice do not display alterations of total KCC2 levels.** Lack of MeCP2 does not induce reductions of KCC2 levels in multiple brain regions. (**A**) Reconstruction of a mouse brain displaying the location of the areas analyzed which do not show KCC2 expression alterations; (**B-C-D-E**) Bar graphs of KCC2 levels (plotted as fold change over WT-Veh) displaying that KCC2 was unaltered in the (B) MOB, (C) AONm, (D) BLA and (E) Hipp CA2/3 in young adult MeCP2 KO mice in comparison to WT controls. On the side, examples of infrared KCC2 labelled coronal sections, areas in green represent the regions of interest quantified in the analysis. MOB = Main Olfactory Bulb (green), AONm = Anterior Olfactory Nucleus, medial part (purple), BLA = Basolateral Amygdala (yellow), CA2/3 = Hippocampus, area CA2/3 (orange). Data expressed as mean ± SEM, single dots represent single observations from 3-4 animals per group, 2-3 coronal planes per brain region, 2-3 sections per coronal plane, left and right hemisphere data pooled together.

### Supplementary Fig. 3

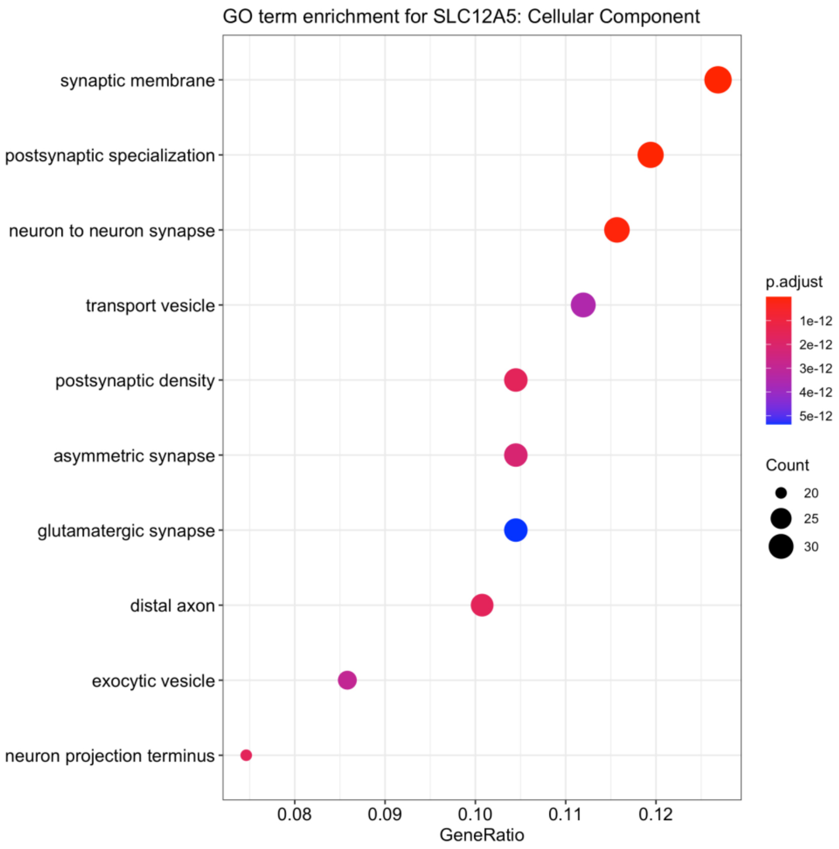


**Supplementary Fig. 3. Gene ontology terms enrichments for Cellular Component (GO:CC) for the genes co-expressed in the *SLC12A5* network.** Graphical representation of GO Cellular Component enrichment for the *SLC12A5* network with FDR adjusted significance levels (color scale) and gene count for each GO category (dot size).

### Supplementary Fig. 4

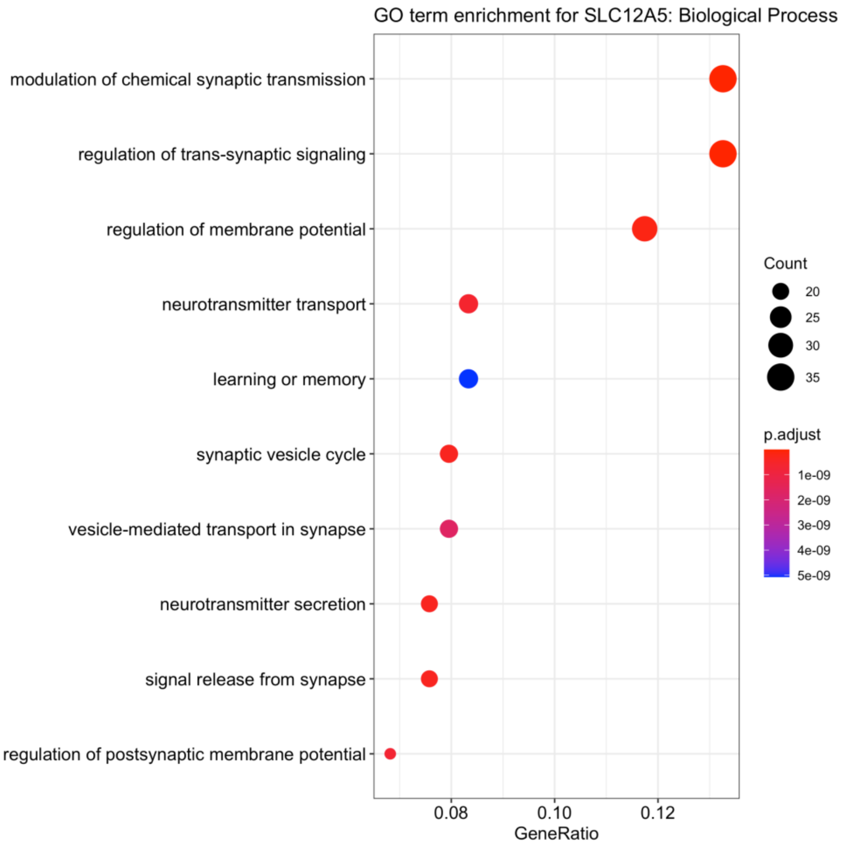


**Supplementary Fig. 4. Gene ontology terms enrichments for Biological Process (GO:BP) for the genes co-expressed in the *SLC12A5* network.** Graphical representation of GO Biological Process enrichment for the *SLC12A5* network with FDR adjusted significance levels (color scale) and gene count for each GO category (dot size).

### Supplementary Fig. 5

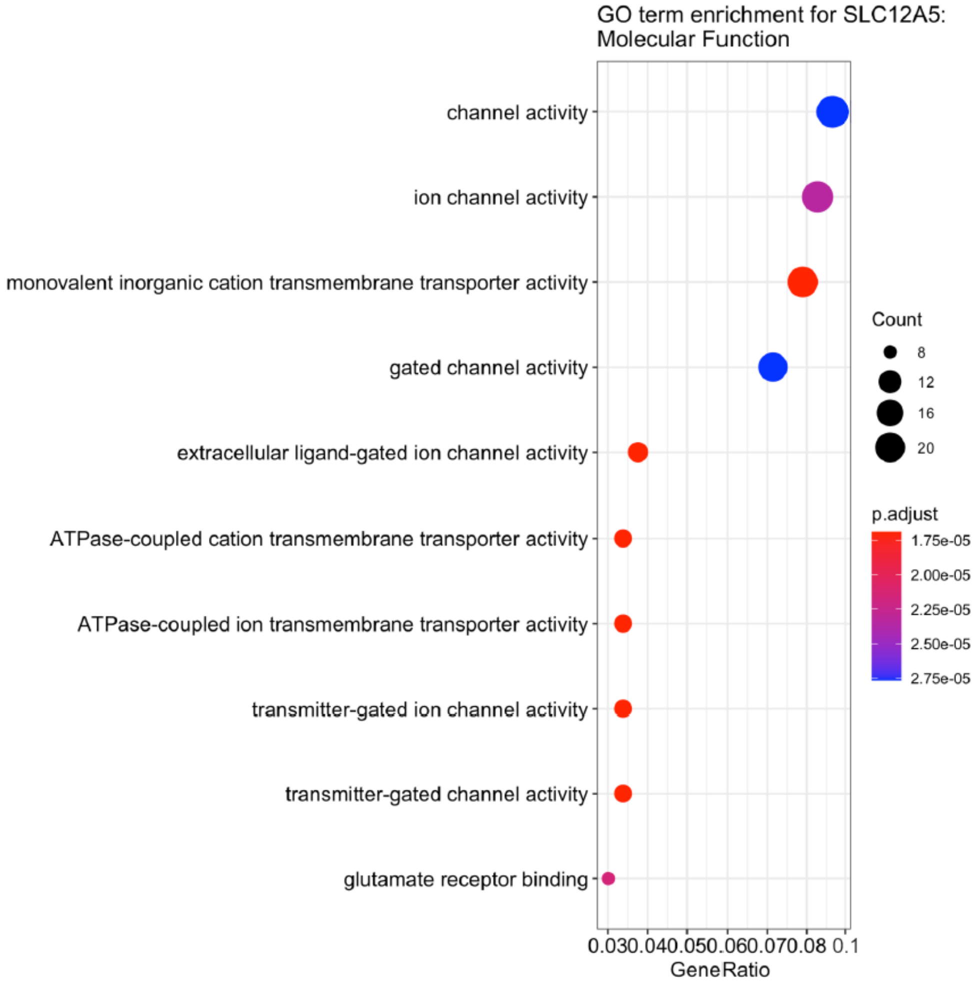


**Supplementary Fig. 5. Gene ontology terms enrichments for Molecular Function (GO:MF) for the genes co-expressed in the *SLC12A5* network.** Graphical representation of GO Molecular Function enrichment for the *SLC12A5* network with FDR adjusted significance levels (color scale) and gene count for each GO category (dot size).

### Bibliography

Abrahams BS, Arking DE, Campbell DB, Mefford HC, Morrow EM, Weiss LA, Menashe I, Wadkins T, Banerjee-Basu S, Packer A. 2013. SFARI Gene 2.0: A community-driven knowledgebase for the autism spectrum disorders (ASDs). *Mol Autism*. 4(1):36.

Hansen KD, Irizarry RA, Wu Z. 2012. Removing technical variability in RNA-seq data using conditional quantile normalization. *Biostatistics*. 13(2):204–216.

Hawrylycz MJ, Lein ES, Guillozet-Bongaarts AL, Shen EH, Ng L, Miller JA, van de Lagemaat LN, Smith KA, Ebbert A, Riley ZL, Abajian C, Beckmann CF, Bernard A, Bertagnolli D, Boe AF, Cartagena PM, Chakravarty MM, Chapin M, Chong J, Dalley RA, Daly BD, Dang C, Datta S, Dee N, Dolbeare TA, Faber V, Feng D, Fowler DR, Goldy J, Gregor BW, Haradon Z, Haynor DR, Hohmann JG, Horvath S, Howard RE, Jeromin A, Jochim JM, Kinnunen M, Lau C, Lazarz ET, Lee C, Lemon TA, Li L, Li Y, Morris JA, Overly CC, Parker PD, Parry SE, Reding M, Royall JJ, Schulkin J, Sequeira PA, Slaughterbeck CR, Smith SC, Sodt AJ, Sunkin SM, Swanson BE, Vawter MP, Williams D, Wohnoutka P, Zielke HR, Geschwind DH, Hof PR, Smith SM, Koch C, Grant SGN, Jones AR. 2012. An anatomically comprehensive atlas of the adult human brain transcriptome. *Nature*. 489(7416):391–399.

Langfelder P, Horvath S. 2008. WGCNA: an R package for weighted correlation network analysis. *BMC Bioinformatics*. 9:559.

Piñero J, Ramírez-Anguita JM, Saüch-Pitarch J, Ronzano F, Centeno E, Sanz F, Furlong LI. 2020. The DisGeNET knowledge platform for disease genomics: 2019 update. *Nucleic Acids Res*. 48(D1):D845–D855.

Raudvere U, Kolberg L, Kuzmin I, Arak T, Adler P, Peterson H, Vilo J. 2019. g:Profiler: a web server for functional enrichment analysis and conversions of gene lists (2019 update). *Nucleic Acids Res*. 47(W1):W191–W198.
